# Multimodal physical evidence uncovers interpretable gene regulatory networks for perturbation prediction

**DOI:** 10.64898/2026.06.05.729520

**Authors:** Zhiwen Yang, Sikai Huang, Ge Bai, Jintong Dong, Jue Wang, Stan Z. Li

**Affiliations:** AI Laboratory, Research Center for Industries of the Future, Westlake University, Hangzhou, China; School of Mathematics, Yangzhou University, Yangzhou, China; College of Computer and Information Science, Fujian Agriculture and Forestry University, Fuzhou, China; School of Future Technology, Dalian University of Technology, Dalian, China; College of Biotechnology and Pharmaceutical Engineering, Nanjing Tech University, Nanjing, China

## Abstract

Gene regulatory networks govern cell fate transitions through dynamic causal mechanisms^1^. Since exhaustively mapping this vast perturbation space experimentally is prohibitive^2^, scalable computational models are essential. Yet, current frameworks fall short because they infer statistical co-expression rather than physical mechanisms, remain blind to non-canonical regulators lacking classical DNA-binding motifs, and fail to generalize across unseen perturbation factors or cell lines. Here we show that a multimodal biophysical framework, VitaGRN, overcomes these barriers by constructing a biophysical regulatory scaffold from multimodal evidence and propagating interactions to capture non-canonical regulators. By leveraging structurally aligned protein embeddings, VitaGRN predicts zero-shot perturbation responses and uncovers non-canonical translational control programs. Notably, VitaGRN demonstrates robust generalization across unseen factors, cell lines, and developmental transitions. Ultimately, VitaGRN generates a confidence-calibrated virtual perturbation atlas spanning over a thousand factors. This resource reframes gene regulatory networks from static correlation graphs into dynamically generalizable and mechanistically transparent models, streamlining wet-lab candidate prioritization.

## INTRODUCTION

In complex biological systems, local interactions among thousands of biomolecules give rise to emergent global phenotypes and cell fate transitions^1^. Gene regulatory networks (GRNs) provide the foundational molecular programs that govern these state changes by orchestrating transcription factor (TF) activity across the genome^3^. Historically, GRN reconstruction has been framed as the mapping of static topological wiring diagrams—delineating pairwise associations between TFs and their target genes. However, the ultimate utility of a regulatory network lies not merely in describing its static topology, but in predicting and manipulating dynamical cellular behaviors under genetic or environmental perturbations. Consequently, the quantitative forecasting of transcriptional responses to perturbations should not be relegated to a downstream application; rather, it must be established as the definitive functional touchstone for validating the accuracy and causal integrity of any inferred GRN.

Unfortunately, existing computational frameworks for GRN inference remain fundamentally constrained under this functional standard, hindered by three distinct, cascading limitations. First, conventional methods rely heavily on statistical co-expression dependencies in single-cell transcriptomic data. Whether utilizing purely correlative metrics^4^ or motif-guided enrichment^5,6^, these frameworks capture statistical associations rather than physical, causal interactions—a limitation systematically confirmed by independent benchmarks^7^. Consequently, a large fraction of inferred edges constitute mechanistic false positives: co-expression arising from shared upstream regulators, cell-cycle co-variation, or technical dropout is statistically indistinguishable from bona fide TF–target binding. As independent benchmarks confirm, existing methods perform only moderately above chance, and the inferred networks are systematically dominated by indirect, length-two associations rather than direct regulatory interactions^7^. Second, to compensate for this correlation noise, traditional methods depend on classic DNA-binding motifs as an exclusive prerequisite. This constraint, however, systematically excludes the vast majority of non-canonical regulators, such as chromatin modifiers, kinases, and translation factors, that lack classical DNA-binding domains^8^. Large-scale single-cell CRISPR perturbation screens^2,9^ now routinely target thousands of genes (including major translational control hubs), yet the overwhelming majority of perturbed factors fall outside the scope of known TF-binding motif databases, leaving them in a systematic computational blind spot for motif-dependent GRN methods. Third, inferred networks suffer from a severe generalization barrier across three distinct axes: unseen perturbation factors, novel cell lines, and dynamic developmental trajectories. Because traditional methods rely on cell-type-specific expression correlation rather than cell-line-invariant biophysical rules, the resulting networks are static blueprints locked to their training context. Consequently, they cannot predict the effects of transcription factors missing from the training data, they cannot transfer to new cell types without complete retraining, and they cannot model dynamic network reorganization along a continuous developmental time-course. In these temporal transitions, transcription factors (such as chromatin rewirers) may drastically rewire their target repertoire without exhibiting large expression changes, a phenomenon entirely invisible to variance-dependent statistical methods. This has driven a deep methodological schism: GRN inference methods^4–6^ produce static, non-transferable networks, while perturbation response predictors^10–13^ achieve quantitative forecasting as black boxes. Neither paradigm delivers a regulatory model that is simultaneously mechanistically explainable and functionally predictive.

To bridge this mechanistic-predictive divide, we must shift the paradigm of GRN construction from statistical pattern matching of expression profiles^7,14^ to models anchored by cell-line-invariant biophysical observations. Because DNA sequence and protein structure remain invariant across different cell types and developmental stages, networks grounded in these physical realities naturally support cross-context generalization. This shift is grounded in a fundamental principle of structural biology: the amino acid sequence of a protein dictates its three-dimensional fold, which in turn determines its function^15,16^. For transcription factors, this implies that a TF’s structure determines its DNA-sequence affinity and target specificity^8,17,18^; structurally similar TFs often exhibit overlapping DNA-binding preferences and partially shared regulatory targets^19^, which can lead to correlated perturbation responses in certain contexts^9,20^. Recent computational advances in structural biology and functional genomics^21,22^ have finally rendered the genome-wide modeling of these biophysical relationships tractable, enabling us to map physical protein folds into functionally aligned representation spaces^23^ and systematically evaluate how genomic sequence variation affects transcription.

To address these challenges, we present VitaGRN, a unified computational framework that bridges genomic regulatory sequence and protein structural homology to reconstruct gene regulatory networks that are simultaneously mechanistically traceable and functionally predictive (**Figure 1**). VitaGRN shifts the paradigm of GRN construction by treating transcriptomic response to perturbations as the definitive functional validator for network inference. We achieve this through a dual-component design consisting of a biophysical regulatory scaffold and a structure-guided response predictor. By utilizing sequence-to-expression mapping and chromatin state signals, the regulatory scaffold constructs physically grounded edge weights, which are propagated through a protein-protein interaction cascade to capture both classical TFs and motif-lacking non-canonical regulators. The response predictor then leverages structurally aligned protein embeddings to project transcription factor structures into a functional representation space, transferring observed perturbation effects to unseen regulators via Bayesian ridge regression. This integration enables zero-shot prediction of transcriptional responses to novel perturbations, translating the static biophysical network into an operational, functionally testable model.

**Figure 1.**
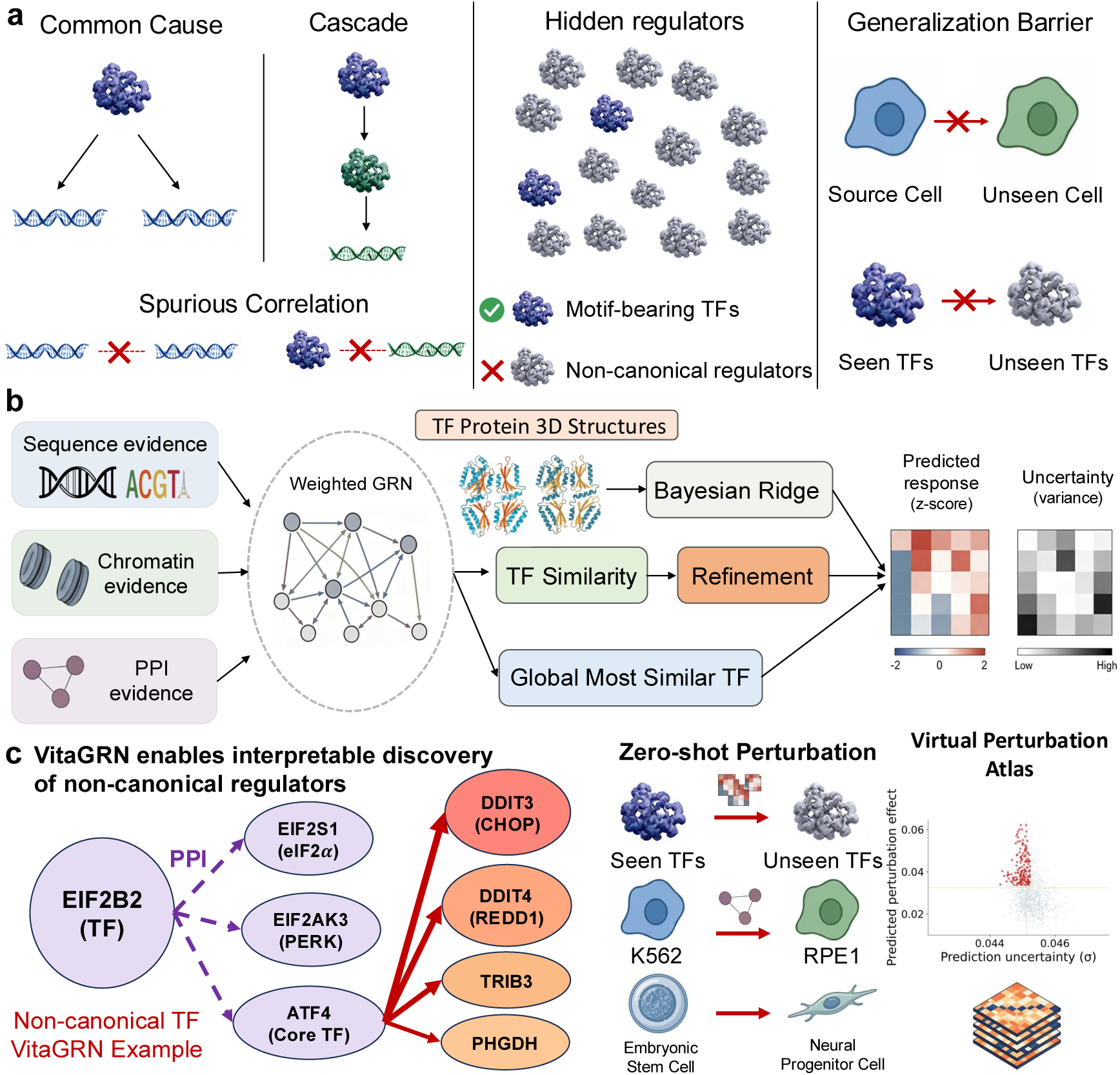
| The VitaGRN framework for mechanism-driven and zero-shot regulatory network inference. **a**, Schematic illustrating three major limitations of current GRN paradigms: false regulatory edges stemming from statistical co-expression noise, the exclusion of non-canonical regulators lacking direct DNA motifs, and strict generalization barriers when transferring models to unseen transcription factors (TFs) or novel target cell types. **b,** Overview of VitaGRN’s dual-engine architecture. The Physics-Grounded Regulatory Scaffold Engine integrates multi-omics data (genomic sequence via in-silico mutagenesis, chromatin accessibility, and protein-protein interaction networks) to con-struct a causal regulatory prior. The Structure Anchor Engine leverages 3D protein structures (ProTrek embeddings) to define functional similarity across TFs. The two engines are integrated via Bayesian ridge regression to achieve end-to-end, confidence-calibrated perturbation predictions. **c,** Summary of breakthrough capabilities and validated discoveries enabled by VitaGRN, including tracing interpretable multi-hop pathways (e.g., the EIF2B2 PPI cascade to ATF4 and stress response genes), executing zero-shot transfer and cross-context generalization (e.g., from leukemia K562 to epithelial RPE1 without retraining), and generating uncertainty-calibrated Virtual Perturbation Atlases for hypothesis prioritization.

We systematically evaluate the utility, generalization, and robustness of VitaGRN across diverse real-world benchmarks and biological systems. First, topological evaluations and direction auditing against a hierarchical gold-standard library in leukemia cells show that VitaGRN recovers causal regulatory wiring with substantially higher accuracy, outperforming existing statistical and motif-based baselines (**Figure 2**). Second, we demonstrate that the framework successfully charts pathways for non-canonical regulators, tracing the integrated stress response cascade from the translation factor EIF2B2 through its physical protein interaction network to downstream target genes (**Figure 3**). Third, we show that the structure-guided response predictor generalizes robustly across large structural distances to predict perturbations of unseen transcription factors, and transfers zero-shot from leukemia cells to normal epithelial cells without retraining (**Figure 4**). Fourth, to overcome the static network barrier, we apply the regulatory scaffold to map continuous dynamic cell-state transitions and construct state-specific networks de novo. We demonstrate that VitaGRN captures temporal regulatory network rewiring and identifies distinct transcription factor modes, including chromatin rewirers, which constitute a hidden regulatory mode invisible to standard correlative approaches, during H9 embryonic stem cell differentiation to neural progenitor cells (**Figure 5**). Finally, we characterize the robustness boundaries of VitaGRN under sequence noise and input sparsity, and generate a global, confidence-calibrated Virtual Perturbation Atlas for candidate prioritization (**Figure 5**).

**Figure 2.**
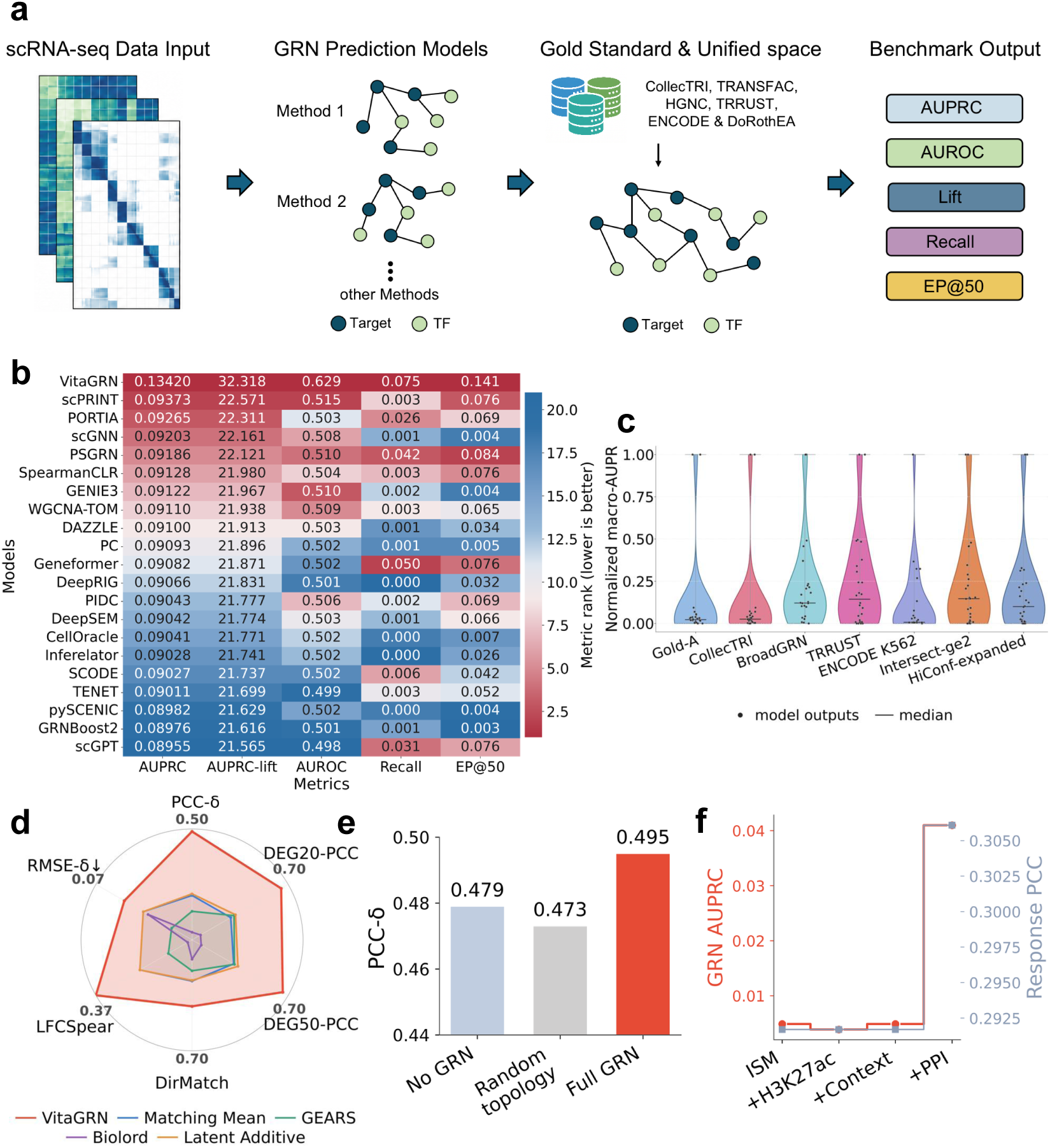
| Biophysical scaffold evaluation and perturbation response prediction in K562. **a**, Unified evaluation schema separating candidate-space definition from gold-standard selection. The schematic illustrates the projection of all model outputs onto a shared directed transcription factor (TF)target edge schema to ensure fair benchmarking. **b,** Rank-colored heatmap of model performance across five metrics comparing VitaGRN to 20 GRN reconstruction methods. Heatmap colors represent normalized ranks (lower is better), with model names alphabetically sorted. **c,** Violin plot showing relative macro-AUPRC distributions across seven independent gold standards for all benchmarked methods, overlaid with individual model data points and median markers. **d,** Radar chart showing normalized ranks of five models across six response prediction metrics, including PCC-*δ*, DEG20-PCC, DEG50-PCC, DirMatch, LFCSpear, and RMSE-*δ* (where lower is better and the axis is inverted to align with other metrics). The outer boundary represents rank 1 (best performance). **e,** Response prediction accuracy (PCC-*δ*) comparing the structure anchor baseline (No GRN), random topology, and the full biophysical scaffold (Full GRN). Data are presented as mean values with error bars representing 95% confidence intervals from bootstrap resampling. **f,** Step curves tracking GRN AUPRC (left axis) and response PCC (right axis) as successive layers of physical evidence (ISM, H3K27ac active chromatin, cell context co-expression, and PPI cascade) are integrated.

**Figure 3.**
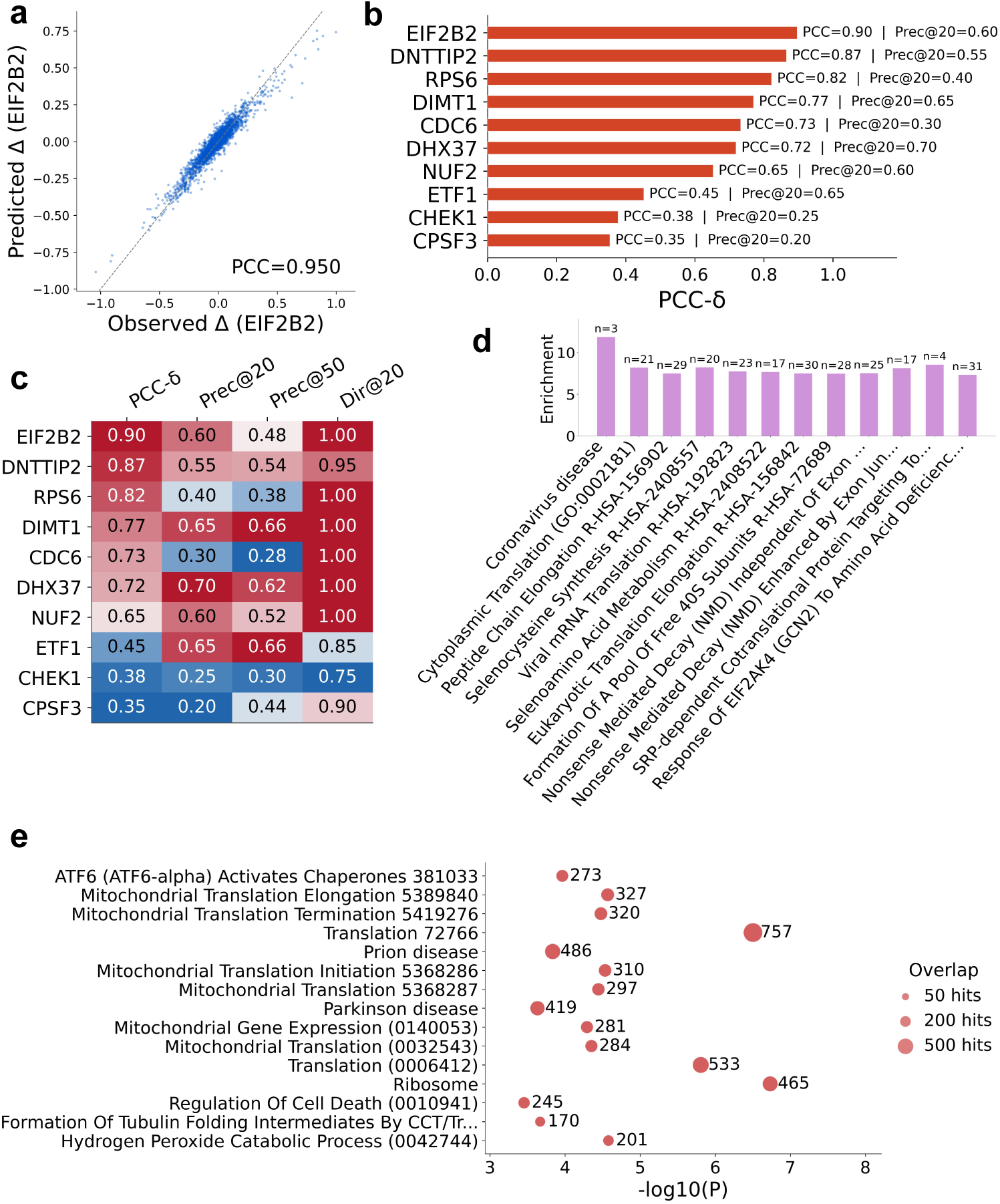
| Predictability and validation of non-canonical regulators through protein interaction cascades. **a**, Genome-wide predicted versus observed differential expression scatter plot for the non-canonical translation initiation factor EIF2B2. The dashed line indicates the y=x identity line, with Pearson correlation coefficient (PCC) annotated. **b,** Bar chart showing prediction accuracy (PCC-*δ*) of the top 10 case-study transcription factors, including non-canonical regulators, annotated with downstream target gene precision (Prec@20). **c,** Multidimensional scorecard heatmap de-composing the prediction performance of the top 10 case-study translation factors across PCC-*δ*, Prec@20, Prec@50, and Direction Accuracy (Dir@20). **d,** GO and Reactome pathway enrichment bar chart of target genes associated with novel predicted edges. Data are presented as mean − log_10_(P-value) across transcription factors, with n indicating the number of distinct regulators targeting each pathway. **e,** Pathway enrichment bubble plot showing statistical significance (− log_10_ P-value) and scale (bubble size representing overlap hits) of biological processes enriched among novel-edge targets.

**Figure 4.**
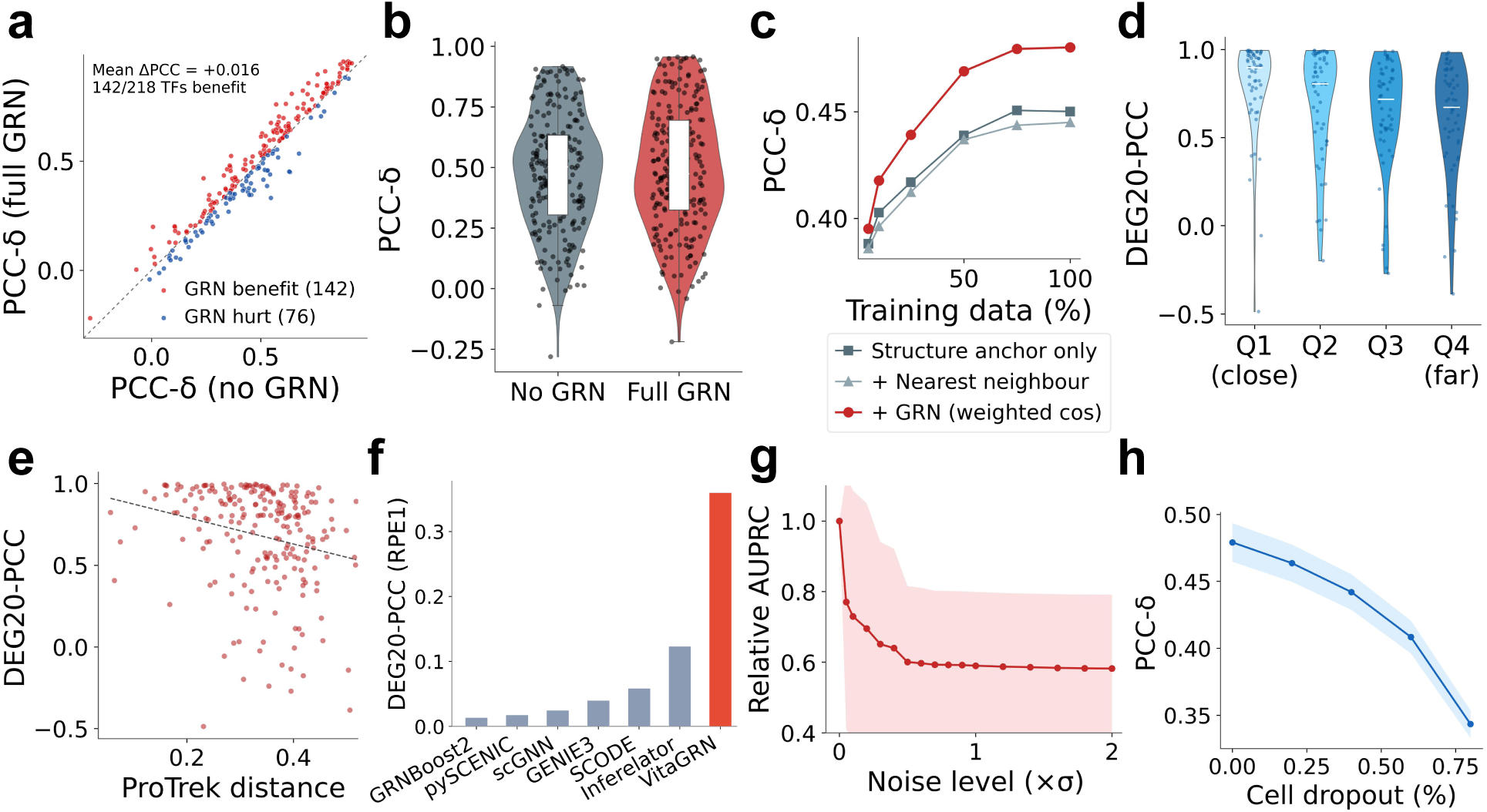
| Generalization across protein-structure distances and cross-cell-line transfer. **a**, Scatter plot of per-TF prediction accuracy (PCC-*δ*) comparing the Full GRN configuration to the structure-anchor-only baseline (No GRN), showing TFs that benefit from or are hurt by topological integration. **b,** Box plot and half-violin distributions comparing overall prediction accuracy (PCC-*δ*) between No GRN and Full GRN configurations across all 218 held-out test TFs. **c,** Data efficiency curves showing prediction accuracy (PCC-*δ*) as a function of training dataset coverage (from 5% to 100%) for three configurations: structure anchor only, nearest-neighbor, and full GRN. **d,** Generalization accuracy (DEG20-PCC) stratified by structural distance quartiles (Q1 to Q4) between the held-out TF and the training set. Medians are indicated by white markers. **e,** Scatter plot illustrating the continuous relationship of prediction accuracy (DEG20-PCC) against structural distance (ProTrek distance) for all test TFs, overlaid with a linear regression fit line (dashed). **f,** Cross-cell-line response transfer accuracy (DEG20-PCC) from leukemia (K562) to normal epithelium (RPE1) comparing VitaGRN to other baseline methods without target-cell retraining. **g,h,** Robustness curves evaluating model stability in terms of relative AUPRC under synthetic sequence-score noise injection (g) and PCC-*δ* under simulated cell dropout sparsity (h). Shaded regions indicate 95% confidence bounds across 5 random seeds.

**Figure 5.**
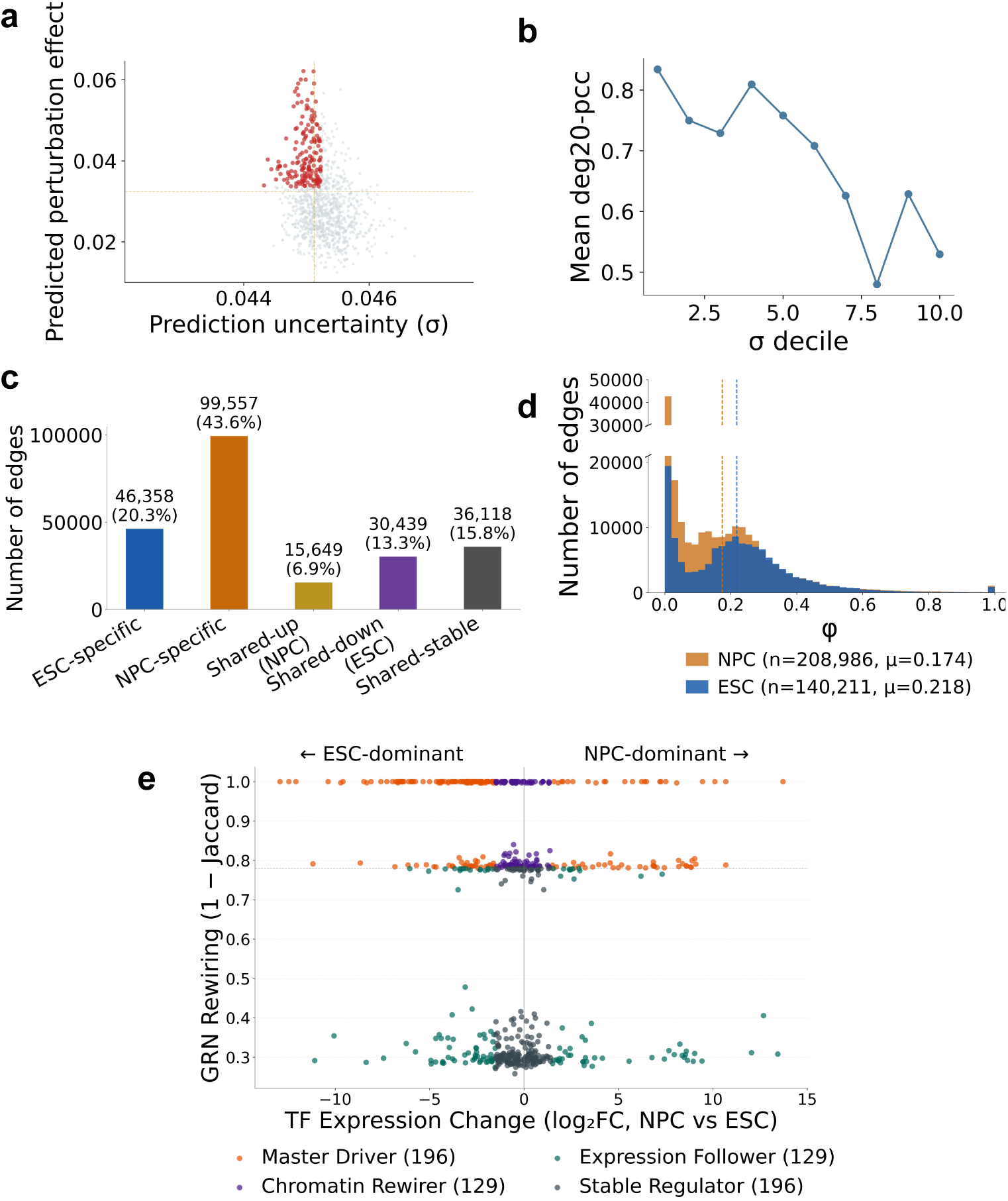
| Virtual perturbation atlas and cell-state regulatory network rewiring. **a**, K562 Virtual Perturbation Atlas scatter plot mapping predicted perturbation effect size magnitude against Bayesian uncertainty (*σ*). The dashed quadrant lines highlight high-effect, low-uncertainty regulatory pairs prioritized for validation. **b,** Uncertainty calibration curve linking predicted uncertainty deciles (*σ* deciles, sorted low to high) with observed prediction accuracy (Mean DEG20-PCC). **c,** Bar chart showing the number and percentage of evaluated (TF, gene) regulatory pairs classified into five categories (ESC-specific, NPC-specific, Shared-up, Shared-down, Shared-stable) across H9 embryonic stem cells (ESCs) and H9-derived neural progenitor cells (NPCs). **d,** Overlapping histograms showing the distribution of non-zero edge weights (*ϕ*) in H9 ESC and H9 NPC states, with dashed lines indicating the respective mean values. **e,** Transcription factor functional regulatory quadrant mapping target gene Jaccard rewiring distance against expression fold change (log_2_ FC, NPC vs. ESC) across 650 factors. Quadrants define Master Drivers, Chromatin Rewirers, Expression Followers, and Stable Regulators.

## RESULTS

### A biophysical framework for inference

GRN inference and perturbation response prediction have historically been treated as separate problems: GRN methods produce static wiring diagrams from expression data, while perturbation predictors map TF identity to transcriptomic outcome without mechanistic interpretability. Traditional GRN paradigms typically suffer from three major limitations: false regulatory edges caused by statistical co-expression, the exclusion of non-canonical regulators lacking DNA-binding motifs, and strict generalization barriers when extrapolating to unseen transcription factors (TFs) or novel cell types (**Figure 1a**). We asked whether both inference and prediction could be unified within a single physically grounded framework. The central output of VitaGRN is a gene regulatory network (GRN) whose edges are anchored in physical observables (genomic sequence, chromatin state, and protein structure) rather than in statistical correlations.

VitaGRN achieves these properties through a two-layer, dual-engine architecture, as detailed in **Figure 1b**. First, the Physics-Grounded Regulatory Scaffold Engine determines potential regulatory paths by integrating multimodal physical evidence. Specifically, it starts from genomic sequence and uses in-silico mutagenesis (ISM) via the AlphaGenome sequence-to-expression model^22^ to estimate sequence-level regulatory potentials. These evidence scores are then calibrated by H3K27ac active-enhancer signals for physical confidence, modulated by cell-type-specific co-expression for functional relevance, and expanded through the protein-protein interaction (PPI) network to non-canonical factors lacking direct motifs. This process outputs a weighted causal GRN and a sparse topology prior (*ϕ*^∼^). Second, the Structure Anchor Engine defines functional similarities between unseen and known TFs using protein structure embeddings from the ProTrek model^23^ derived from 3D protein structures. Finally, a central integration hub combines the prior from the scaffold engine with the functional similarities from the structure engine via Bayesian ridge regression, generating end-to-end, confidence-calibrated perturbation predictions.

Critically, this dual-engine design serves as a functional validation of the GRN: the biophysical scaffold supplies a prior on *which genes are likely to respond*, and the structure anchor quantifies *by how much*. If the inferred GRN carries real regulatory mechanisms rather than spurious correlations, it should significantly improve perturbation prediction beyond what sequence or structure alone can provide.

We systematically evaluated VitaGRN across diverse biological contexts. We anchored our core functional validation on the large-scale K562 Perturb-seq dataset^2^, evaluating both the quality of the reconstructed physical GRN and the accuracy of zero-shot perturbation response predictions. To rigorously test the generalizability of our framework, we evaluated cross-cell-line transferability using RPE1 Perturb-seq data^2^ without any cell-specific model retraining. Detailed dataset specifications and the evaluation pipeline are provided in the Methods.

As illustrated in **Figure 1c**, these extensive evaluations demonstrate several robust capabilities and validated discoveries of the VitaGRN framework. First, it uncovers interpretable pathways for non-canonical regulators; for example, it successfully traces the physical regulation path from the non-canonical translation factor EIF2B2 through its PPI cascade to the core TF ATF4, revealing its direct transcriptional activation of integrated stress response (ISR) genes. Second, the physics-grounded formulation enables accurate zero-shot perturbation transfer to unseen TFs. Third, it achieves seamless cross-context generalization, transferring predictions from leukemia cells (K562) to normal epithelium (RPE1) without retraining. Finally, the framework generates a confidence-calibrated Virtual Perturbation Atlas (Δ) with explicit uncertainty bands, establishing a reliable and traceable resource for downstream hypothesis generation.

### Physical edges reduce correlation noise

K562 is a chronic myeloid leukemia (CML) cell line, widely used as a model for hematopoietic gene regulation and the largest source of genome-wide Perturb-seq data^2^. We constructed VitaGRN’s biophysical regulatory scaffold on K562 and first asked whether the resulting GRN topology is competitive under a unified matched-space evaluation. Existing GRN benchmarks are difficult to compare because methods are evaluated on different gene universes, edge candidates, and gold standards. To this end, we evaluated VitaGRN in a K562 GRN benchmark that separates candidate-space definition from gold-standard selection (**Figure 2a**). The benchmark projects all model outputs onto a shared directed transcription factor (TF)–target edge schema, establishing a unified evaluation space on K562 (the main candidate space, S_main_) and evaluating them against the same hierarchical reference library (the integrated reference standard, G_ref_) under a unified fixed-space evaluation (**Extended Data Fig. 1**).

The unified evaluation showed that VitaGRN ranked first among the 21 evaluated models, achieving a macro-AUROC of 0.6286 (**Figure 2b**). Crucially, while traditional statistical and deep-learning inference methods collapsed to near-random performance (AUROC ≈ 0.50) due to their inability to distinguish direct causal binding from indirect co-expression noise, VitaGRN maintained high causal accuracy. This demonstrates that anchoring edges in physical observables successfully filters out the massive statistical false positives that plague conventional inference. Furthermore, VitaGRN exhibited high early-budget precision and an 80.2% edge sign consistency, proving that its biophysical edge weights natively capture excitatory and inhibitory logic (Supplementary Fig. 2; Supplementary Table 1). This topological superiority held robustly across seven independent gold-standard definitions, significantly outperforming the second-best method (one-sided Wilcoxon signed-rank test, *P <* 2.2 × 10^−16^; **Figure 2c**), with detailed rank structures across all candidate spaces and gold standards provided in the supplementary evaluation rank heatmap (**Extended Data Fig. 2**).

Having established that the scaffold produces high-quality causal topology, we evaluated whether VitaGRN’s end-to-end predictions agree with experimentally observed perturbation responses. Rathe than just reporting overall correlation, we audited three complementary dimensions: target-gene recovery, direction accuracy, and effect-size correlation (**Figure 2d**). VitaGRN accurately predicted regulatory directions for all test TFs, achieving an overall direction accuracy of 0.862 (95% CI: [0.851, 0.873]), with confidence bounds strictly superior to all baselines (Supplementary Table 2 and Supplementary Note 1). Target-gene recovery and effect-size correlation similarly exceeded all comparators, confirming that end-to-end learning successfully translates GRN topology into quantitative response magnitudes, with prediction quality strongly correlating with the biological perturbation effect magnitude (**Extended Data Fig. 3**). Crucially, this advantage was not driven by differential TF coverage, as VitaGRN remained superior even on a strict common-ground subset of TFs evaluable across all methods. Finally, an ablation study confirmed that replacing the weighted scaffold with a random, unweighted, or structure-only topology severely degraded prediction accuracy (**Figure 2e**). This establishes a perfect causal loop: high-quality physical topology is a necessary prerequisite for accurate quantitative predictions.

### Resolving non-canonical regulators

A cumulative ablation confirmed that each layer of the biophysical scaffold, consisting of sequence potential, epigenetic gating, and single-cell co-expression, progressively enhanced GRN quality. Crucially, propagating these weights through the protein-protein interaction (PPI) cascade extended coverage from 13 motif-possessing TFs to hundreds of non-canonical factors. This addition of PPI caused GRN AUPRC and downstream response prediction accuracy to rise in synchrony, proving that physical interaction cascades successfully link primary sequence signals to broad perturbation responses.

VitaGRN’s novel-edge perturbation validation rate yielded a 78.2× higher validation ratio than expected by random chance (Fisher’s exact test, *P* = 4.1 × 10^−42^), substantially exceeding all baselines (Supplementary Fig. 4). These biologically validated novel edges predominantly connect non-canonical factors, such as kinases, epigenetic modifiers, and RNA processing factors, that lack classical DNA-binding motifs and are therefore completely invisible to traditional motif-based GRN methods.

To illustrate the interpretability afforded by the GRN, we examined local biological case studies. EIF2B2 was the strongest case (**Figure 3a**): PCC-*δ* = 0.950, with 12 of its top-20 DEGs covered by the VitaGRN local network and 100% direction consistency. The covered genes include well-established ISR effectors downstream of the ATF4–CHOP axis—namely DDIT3 (CHOP), DDIT4 (REDD1), and TRIB3—alongside PHGDH (linked to ATF4-associated serine biosynthesis) and the chaperone HSPA8^24^. Biologically, EIF2B2 encodes a subunit of the eIF2B translation initiation complex. Its genetic attenuation triggers global translation inhibition while selectively enhancing the translation of the core stress responder ATF4^25^ (detailed biochemical mechanisms and literature support in Supplementary Note 8). Once translated, ATF4 directly activates down-stream transcriptional effectors (DDIT3, DDIT4, and TRIB3)^26^. This regulatory chain is indirect: EIF2B2 is a translation factor that does not bind DNA, and its transcriptional effects are mediated entirely through ATF4 translational control. VitaGRN captures this indirect regulation because the PPI cascade propagates evidence from motif-bearing TFs through the translation machinery to ISR effector genes, converting a physical interaction network into a mechanistic regulatory hypothesis.

Comparing EIF2B2 with two ribosomal proteins, RPS6 (PCC-*δ* = 0.864, 40S subunit) and RPL13 (PCC-*δ* = 0.957, 60S subunit), reveals a broader pattern. All three achieve strong prediction performance and converge on the translation apparatus, yet their downstream targets differ (**Extended Data Fig. 4**): EIF2B2 uniquely affects ISR-specific genes (DDIT3, DDIT4, TRIB3) via the ATF4 relay, whereas RPS6 and RPL13 predominantly perturb ribosomal protein and nucleolar stress genes^27,28^ (**Figure 3b, c**). This illustrates that VitaGRN does not merely learn a generic “translation stress” signature but distinguishes the distinct signalling conduits, including ISR, nucleolar stress, and translational feedback, through which different transcription factors exert their regulatory effects.

Additional cases spanning translation, mitosis, RNA processing, and metabolism are documented in **Figure 3b, c**. At the pathway level, GO enrichment of novel-edge targets confirms that VitaGRN’s predictions converge on biologically coherent processes, with ribosomal and translation pathways dominating the top enriched terms (**Figure 3d, e**). These case studies demonstrate that VitaGRN successfully traces physical evidence chains to uncover hidden regulatory hubs.

### Zero-shot prediction and cell transfer

A central premise of VitaGRN is that protein structure is informative of regulatory function: structurally similar TFs often exhibit overlapping DNA-binding preferences and may, in specific contexts, regulate overlapping target sets; in perturbation screens, related regulators can also produce correlated transcriptional response patterns. We first asked whether structurally aligned protein embeddings capture this biological principle. The UMAP projection of all 1,087 TF embeddings (**Extended Data Figure 5a**) reveals that TFs cluster by functional category: motif-possessing TFs (brick-red, 14 TFs) form a distinct, compact neighbourhood separate from the broader distribution of non-canonical factors (gray, 1,073 TFs). This structural organisation provides the biological foundation for zero-shot transfer: if a held-out TF is structurally close to training TFs, its response can be inferred from those neighbours.

If the GRN constructed by VitaGRN carries genuine regulatory information, rather than merely fitting statistical patterns, then it should further improve prediction beyond what protein structure alone provides. We tested this proposition on the K562 benchmark, treating 218 TFs as a completely held-out test set and evaluating genome-wide expression response prediction at the perturbation level. The GRN-backed model led comprehensively across all core metrics, achieving a DEG20-PCC of 0.681 (vs. GEARS 0.553; paired t-test, *P* = 3.8 × 10^−12^) and a PCC-*δ* of 0.495 (vs. Matching Mean 0.370; paired t-test, *P <* 2.2 × 10^−16^; see Supplementary Table 2 for the full benchmark results across all 6 evaluation metrics). To directly test whether the GRN topology contributes functional information, we compared prediction accuracy (PCC-*δ*) of the Full GRN configuration to the structure-anchor-only baseline (No GRN) across individual TFs (**Figure 4a**) and their overall distributions (**Figure 4b**). Replacing the real GRN with randomly shuffled TF-target relationships degraded performance, confirming that topological noise harms prediction. Data efficiency analysis as a function of training dataset coverage from 5% to 100% (**Figure 4c**) further showed that the structure anchor alone (with or without nearest-neighbor search) already constitutes a strong baseline at very low training coverage, and the gain from GRN guidance is small but directionally stable, indicating that VitaGRN’s advantage lies primarily in structurally constrained zero-shot extrapolation.

We next tested whether the GRN-backed response predictor generalizes beyond the training distribution across the protein-structure space. We divided the 218 test TFs into four groups by structural cosine distance quartiles (Q1 to Q4; **Figure 4d**). The violin distributions reveal a clear monotonic trend: Q1 (closest structural neighbours) achieved a mean DEG20-PCC of 0.797, Q2 of 0.692, Q3 of 0.637, and Q4 (farthest) retained 0.596. The Q4/Q1 retention ratio of 74.8% for DEG20-PCC demonstrates genuine generalization rather than memorization of nearest-neighbor responses. Notably, even Q4 TFs, whose closest structural neighbours in the training set are substantially different, retain DEG20-PCC well above random, confirming that the structure anchor captures broad functional similarity beyond nearest-neighbour lookup. To analyze this relationship as a continuous gradient rather than discrete quartiles, we plotted prediction accuracy (DEG20-PCC) against structural distance (ProTrek distance) for all test TFs in a joint scatter plot with marginal histograms, showing the continuous trend where model performance decays smoothly with increasing physical distance as outlined by the black dashed fitted line (**Figure 4e**).

Finally, we tested whether the physical regulatory scaffold itself, being built from cell-line-invariant genomic sequence and protein structure, transfers across cell lines. We transferred the K562-constructed scaffold directly to RPE1, a non-cancerous retinal pigment epithelial cell line, without any RPE1-specific retraining (**Figure 4f**). VitaGRN end-to-end predictions achieved DEG20 PCC = 0.360 and PCC-*δ* = 0.120 (47 shared TFs), clearly exceeding all GRN-only first-order propagation baselines (highest ∼0.06). GRN-only methods produced cross-cell-line DEG20-PCC values near zero, reflecting the inability of pure topological propagation to transfer effectively in a zero-shot cross-cell-line setting. These two capabilities are distinct but mutually reinforcing: the response predictor generalizes across protein-structure distance, while the physical regulatory scaffold enables cross-cell-line transfer without retraining.

Robustness analyses confirmed that VitaGRN’s predictions remain stable under typical sequence-level noise and severe single-cell dropout sparsity (Supplementary Fig. 8; **Figure 4g, h**).

### Application boundaries and cell states

We designed robustness and boundary analyses to determine when VitaGRN should be expected to generalize. Extensive structural ablations, including evaluations of protein structural embedding validity and pLDDT score confidence (**Extended Data Fig. 6**), along with task-level transfer boundaries under chemical perturbations (**Extended Data Fig. 7**), confirm the robustness of VitaGRN’s structural anchor and define its application boundaries (Supplementary Figs. 9 and 10; Supplementary Table 3).

Virtual perturbation atlas. VitaGRN generated a global virtual perturbation atlas for 1,087 K562 perturbation factors across 5,000 genes, complete with Bayesian uncertainty estimates, in just 34 seconds (**Figure 5a**, with the overall uncertainty distribution shown in **Extended Data Fig. 8**). Crucially, these uncertainty estimates display a strict monotonic relationship with observed prediction accuracy (**Figure 5b**). When the model predicts low uncertainty, the prediction is highly accurate (PCC *>* 0.8), providing a reliable confidence calibration metric to prioritize candidates for wet-lab validation.

Having established the reliability and boundaries of VitaGRN in static cell lines, we next evaluated its ability to overcome the static network limitation by capturing continuous dynamic regulatory rewiring during developmental cell fate transitions. **Regulatory network rewiring during H9 ESC-to-NPC differentiation.** To demonstrate the applicability of VitaGRN’s regulatory scaffold to mapping temporal cell fate transitions, we applied it to H9 embryonic stem cells (ESCs) and H9-derived neural progenitor cells (NPCs) independently, using state-specific RNA-seq from an H9 ESC-to-NPC differentiation dataset in which PAX6 marks the neuroectoderm transition^29^. For each state, 500 highly expressed genes were scored against 640 (ESC) / 617 (NPC) motif-matched TFs through the full pipeline: ISM scoring, DNase gating, and Spearman co-expression modulation. After aggregating to (TF, gene) pairs, the union space comprised 303,769 regulatory pairs.

Edge-weight validation and state classification. Classifying all 303,769 regulatory pairs revealed that NPC-specific edges significantly outnumbered ESC-specific edges (**Figure 5c**), consistent with the massive acquisition of new transcriptional programs during neural lineage commitment. Furthermore, edge weights (*ϕ*) successfully captured state-specific regulatory relationships (**Figure 5d**), with top-tier edges showing over twofold higher validation rates against orthogonal curated databases like CollecTRI (Fisher’s exact test, *P* = 1.3 × 10^−8^; Supplementary Fig. 12; Supplementary Table 4).

Transcription factor regulatory mode classification. By combining TF expression change (log_2_ fold-change of D18 NPC vs. D0 ESC, where positive values denote NPC-upregulation and negative values denote ESC-upregulation) with GRN rewiring magnitude (Jaccard distance between ESC and NPC target gene sets), we classified 650 TFs into four functional categories (**Figure 5e**). First, *Master Drivers* (196 TFs) are characterized by large expression changes combined with complete GRN rewiring; ESC-specific examples include OVOL2 (770× downregulated, Jaccard = 0) and IRF6 (1,310× downregulated), while NPC-specific examples include TBR1 (1,640× up-regulated) and DMRTA2 (1,100× upregulated). Second, *Chromatin Rewirers* (129 TFs) display near-stable expression but complete GRN reorganization, representing a shift in regulatory target repertoire without changes in abundance; TP63 typifies this class with only a 1.5× expression change but Jaccard = 0 between states, yet achieving the highest mean edge weight among all 650 TFs (*ϕ*^ˉ^_NPC_ = 0.788). Third, *Expression Followers* (129 TFs) show large expression changes but maintain highly stable target gene repertoires. Fourth, *Stable Regulators* (196 TFs) display both constitutive expression levels and stable target gene sets. The identification of Chromatin Rewirers is uniquely enabled by the regulatory scaffold’s ability to predict edges independently of TF–target co-expression patterns. Correlation-based GRN methods cannot detect TFs whose regulatory targets change without accompanying expression changes.

Candidate prioritization for validation. We ranked all TFs by a composite score integrating rewiring magnitude, edge confidence, state specificity, and biological novelty. The top candidates span both Master Driver and Chromatin Rewirer categories and have established or potential disease relevance (**Supplementary Table 5**; detailed validation protocols in Supplementary Methods).

Biological implications of state-specific networks. This analysis demonstrates that the regulatory scaffold, built without perturbation training data, can (1) construct cell-state-specific GRNs de novo, (2) capture biologically coherent regulatory reorganization during differentiation, and (3) identify specific, testable hypotheses about TF-level rewiring—including Chromatin Rewirers that are invisible to correlation-based methods. The approach is generalizable to any cell state with matched AlphaGenome ontology tracks and RNA-seq data.

## DISCUSSION

The use of multimodal physical evidence fundamentally distinguishes VitaGRN from conventional methodologies, shifting gene regulatory network (GRN) inference from statistical coexpressio to physical causation. Conventional expression-based methods inevitably struggle with correlation noise and are fundamentally incapable of modeling the activity of non-canonical regulators that lack classical DNA-binding motifs. By integrating an in-silico mutagenic sequence prior with protein-protein interaction (PPI) cascades, VitaGRN constructs a biologically traceable regulatory scaffold. This enables the precise quantification of activity for hidden regulatory hubs, such as translation initiation factors mediating stress responses, resolving a major blind spot in current single-cell network inference. Furthermore, unlike purely predictive deep learning frameworks, VitaGRN provides an explicit mechanistic provenance for every predicted perturbation response, ensuring that computational predictions can be directly translated into interpretable wet-lab hypotheses.

Specifically, VitaGRN fundamentally extends upon previous linear models, such as CellOracle, by integrating protein and sequence foundational models like ProTrek and AlphaGenome. Previously, constructing GRNs often required the fraught alignment of scRNA-seq and ATAC-seq data, a process inherently plagued by dropouts, batch effects, and pervasive noise. By contrast, VitaGRN bypasses these multi-omics alignment artifacts by constructing a robust physical skeleton derived directly from AlphaGenome’s physical data, requiring only DNA sequence as input. Furthermore, while most previous methods fail to generalize to “unseen” perturbations or non-classical regulators, we leverage protein interaction priors to bridge this gap. Ultimately, our framework represents a principled synergy of multiple orthogonal data sources—demonstrating that true predictive power stems from mechanistic integration rather than naively scaling scRNA data or merely augmenting training sets with more perturbation data to brute-force model accuracy.

A defining capability of VitaGRN is its ability to overcome the generalization barriers that severely limit statistical models. Because the foundational layers of the regulatory scaffold, comprising DNA sequence regulatory potential and protein 3D structure embeddings, are invariant across cell types, they form a universal and transferable prior. We demonstrated that this physical invariance enables direct zero-shot cross-cell-line transfer without any target-cell retraining. More importantly, it successfully captures dynamic cell-state-specific network rewiring during continuous developmental trajectories, identifying critical “Chromatin Rewirers” that undergo complete target reorganization despite maintaining stable expression levels. Such dynamic regulatory modes are fundamentally invisible to classical correlation-based methods that depend entirely on expression variance, highlighting how structurally anchored priors unlock the discovery of elusive developmental regulators.

VitaGRN also offers notable computational advantages for large-scale analysis. By leveraging pre-computed foundation model embeddings and efficient Bayesian regression, the framework generated a comprehensive virtual perturbation atlas encompassing 1,087 transcription factors across 5,000 genes, complete with precise uncertainty calibration bounds, in under a minute. However, there are certain limitations associated with the current framework. First, the response predictor relies on a linear architecture; it cannot capture complex nonlinear regulatory dynamics such as combinatorial epistasis or transcriptomic saturation effects. Second, the reliance on high-throughput PPI networks inherently introduces interaction false positives that require strict thresholding. Third, VitaGRN’s predictive validity is strictly confined to genetic perturbations; extrapolating this framework to chemical or small-molecule drug responses remains an unresolved challenge. Fourth, because the framework relies heavily on the DNA sequence to construct its physical skeleton, it may be inherently limited in capturing purely epigenetic silencing or variations independent of sequence changes. Finally, the framework’s overall accuracy is fundamentally bounded by the resolution and reliability of the underlying protein structure and sequence-to-expression foundation models. Crucially, prediction quality is robustly positively correlated with the actual biological perturbation effect magnitude (Spearman *ρ* = 0.41; Extended Data Fig. 3a). While large-effect TFs, predominantly translation and ribosomal components, achieve the highest accuracy and maximum GRN-driven performance gains (Extended Data Fig. 3a, b), medium-effect TFs representing signaling regulators whose subtle effects distribute across downstream cascades remain the most challenging cohort. This highlights the inherent difficulty of capturing attenuated signal transduction and points toward a critical direction for future model refinement.

By constructing physically grounded, highly generalizable, and confidence-calibrated gene regulatory networks, VitaGRN bridges the historical divide between static GRN inference and dynamic perturbation prediction. The framework enables the use of robust statistical techniques to prioritize high-confidence regulatory pairs before committing to costly experimental validation. We anticipate that as large-scale perturbation atlases and structural models continue to evolve, physically constrained frameworks like VitaGRN will be increasingly utilized not only to characterize the molecular mechanisms driving cellular phenotypes but also to investigate a wide range of complex questions in precision medicine and therapeutic target discovery. Finally, while VitaGRN successfully captures direct mechanistic links, the field has not yet fully unraveled the complex emergent effects of global gene regulation. Understanding how localized perturbations cascade into system-wide emergent properties represents a critical frontier for our future research.

## Supporting information

Source Data for Figure 1

Source Data for Figure 2

Source Data for Figure 3

Source Data for Figure 4

Source Data for Figure 5

Source Data for Figure 6

Source Data for Figure 7

Source Data for Figure 8

## METHODS

### Implementation details of VitaGRN

VitaGRN’s regulatory scaffold integrated four layers of physical and functional evidence in a cascaded pipeline: sequence-level *in silico* mutagenesis (ISM), chromatin-state confidence calibration, cell-type-specific co-expression context, and protein–protein interaction (PPI) cascade propagation. The pipeline operated on a predefined gene set; for each gene, the dominant transcription start site (TSS) was selected by querying the AlphaGenome model^22^—a sequence-to-expression predictor that accepts DNA sequence and a cell-type ontology term as input and predicts RNA expression, DNase accessibility, and Micro-C contact maps in a cell-type-aware manner.

#### Motif scanning and candidate filtering

For each gene, we scanned a 200 kb window centered on its dominant transcription start site (TSS) using the JASPAR CORE 2024^30^ and HOCOMOCO v11^31^ position weight matrix (PWM) libraries. Candidate TF binding sites were retained above a prespecified PSSM log-odds score threshold. TFs whose cognate genes were not detectably expressed in the target cell type were removed at this stage. Hierarchical gating was then applied to select high-confidence distal candidates: first, genomic regions were restricted to open chromatin via DNase accessibility gating; second, candidate enhancers were filtered by Micro-C distal contact frequencies using an adaptive contact threshold; finally, an adaptive Top-*N* truncation was applied, retaining the top *N*_typ_ distal candidates for typical TFs and *N*_glob_ for global regulators (*N*_typ_ *< N*_glob_). For cell types lacking Micro-C training data in AlphaGenome, distal candidates are instead prioritized by chromatin accessibility, TF expression level, and genomic distance to the target promoter. All threshold values are reported in Supplementary Table 6.

#### ISM layer

We utilized the AlphaGenome sequence-to-expression model^22^ to estimate sequence-level regulatory potentials by predicting RNA expression from genomic sequence. For each candidate TF motif site, we performed local in-silico mutagenesis (ISM) by predicting the RNA expression readout difference between the wild-type and mutant sequences, where the mutation was introduced via dinucleotide shuffling within the motif footprint. To prevent expression magnitude from dominating the edge weights directly, the raw difference was normalized by the reference expression level of the target gene:

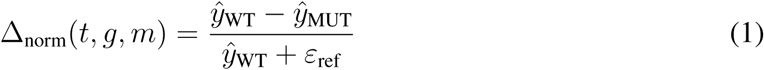

where *ε*_ref_ is a stabilization term. When multiple binding sites existed for the same TF-target pair, we retained the site with the largest absolute effect:

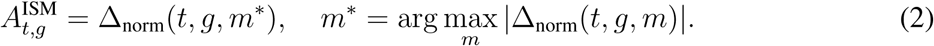

#### Chromatin calibration layer

ISM scores were subsequently calibrated for physical confidence using H3K27ac active-enhancer signals:

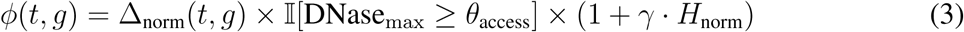

where *H*_norm_ denotes the normalized H3K27ac signal, *θ*_access_ is the minimum DNase accessibility threshold, and *γ* is the chromatin-calibration scaling weight. This step preserved the direction information from sequence evidence while suppressing sites that lacked active chromatin support.

#### Context layer

To introduce cell-type specificity, we estimated TF-target co-expression *C*(*t, g*) from control single-cell profiles using the Spearman correlation coefficient and used only its positive component to modulate the physical evidence:

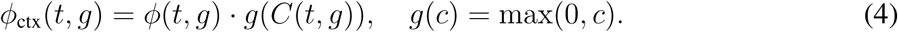

Co-expression was computed from control (unperturbed) single-cell RNA-seq profiles of the target cell type, ensuring that the modulation reflects native regulatory coupling rather than perturbation-induced correlations.

#### PPI cascade layer

For perturbation factors that lacked direct motif evidence, we propagated the regulatory skeleton from ISM-evidenced neighbor TFs using the high-confidence protein–protein interaction network from STRING v12^32^, filtered by a combined interaction score threshold. Regulatory weights were propagated via multi-hop neighbor connections. The final regulatory scaffold was normalized per TF as:

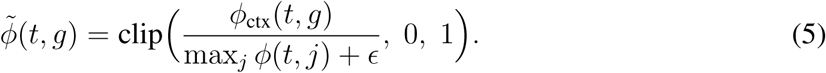

This PPI cascade step enabled non-canonical regulatory factors to acquire traceable functional neighbors and candidate target gene sets.

VitaGRN’s response predictor operates through a single-branch architecture combining three complementary components: a protein-structure-anchored global prior (Structure Anchor), a regulato scaffold-guided cross-TF transfer, and a genome-wide nearest-neighbor supplement. The Structure Anchor provides the primary prediction for every query TF by mapping protein structure embeddings to perturbation response space via Bayesian ridge regression. Cross-TF transfer then applies a mechanism-driven local correction: for genes with physical scaffold support, it blends the Structure Anchor prediction with the observed responses of functionally similar training TFs, where similarity is measured by weighted cosine similarity between GRN edge weight vectors. Finally, the nearest-neighbor supplement provides a genome-wide phenotypic anchor by matching the structure-anchored prediction to the most correlated training TF’s response profile across all genes. The PPI-enhanced regulatory scaffold ensures that 98.4% of perturbation factors (1,060/1,077) have GRN coverage, enabling cross-TF transfer for the vast majority of query TFs. All hyperparameters were selected on the held-out validation set; values are reported in Supplementary Table 6.

#### Structure anchor definition

We utilized structurally aligned protein embeddings from the ProTrek protein structure foundation model^23^ as a global prior. The 1,024-dimensional structurally aligned protein embeddings map protein sequence and structural information into functionally aligned representation spaces^19,20^. Structurally similar TFs often exhibit overlapping DNA-binding preferences and may, in specific contexts, regulate overlapping target sets; in perturbation screens, related regulators can also produce correlated transcriptional response patterns. Given the mean response matrix of *P* training perturbations **Y** ∈ ℝ*^P^* ^×^*^G^* over *G* highly variable genes, we first applied truncated singular value decomposition (SVD) to obtain a low-rank response dictionary **D** ∈ ℝ*^G^*^×^*^K^* with *K* latent components. The SVD coefficients for all training TFs are collected in **C**_train_ ∈ ℝ*^P^* ^×^*^K^*. Let **E**_train_ ∈ ℝ*^P^* ^×^*^d^* be the row-concatenated protein structure embeddings of the training TFs, where *d* is the embedding dimension. We learned the linear mapping from embedding space to SVD coefficient space via ridge regression:

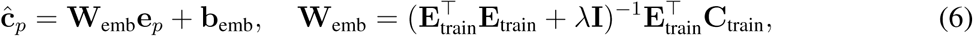

where **W**_emb_ ∈ ℝ^1024×20^, **b**_emb_ ∈ ℝ^20^, and *λ* is the L2 regularization strength. The Bayesian ridge variant (implemented via BayesianRidge from scikit-learn) places a Gamma prior on *λ*, yielding predicted coefficients **c**^_query_ that are decoded to obtain the structure-anchored prediction *δ*^^^_anchor_ = Dc^_query_ ∈ ℝ^5000^. The per-gene uncertainty estimates ***σ*** ∈ ℝ^5000^ are derived from the posterior predictive variance:

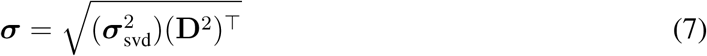

swhere ***σ***^2^ is the SVD coefficient uncertainty vector.

#### Cross-TF transfer learning

The regulatory scaffold connected unseen TFs to training TFs through shared target genes, providing a mechanism-driven local correction. For an unseen TF *t* and a training TF *s*, we defined their functional similarity *S*(*t, s*) via weighted cosine similarity between their regulatory scaffold edge weight vectors ***ϕ***^∼^*_t_, **ϕ***^∼^*_s_* ∈ ℝ^5000^:

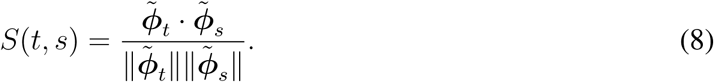

This formulation leverages the full edge weight vectors, preserving the quantitative regulatory-strength information. We selected the top-*K_N_* most similar training TFs as functional neighbors N (*t*). The cross-TF transfer operates exclusively on the *local regulon*, defined as the set of target genes for which the regulatory scaffold provides physical regulatory support. For a gene *g* in the shared regulon (indicated by the binary mask 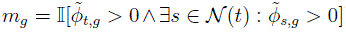, the known responses of the neighbor TFs were blended with the structure-anchored prediction *δ*^^^_anchor,*g*_ using a mixing weight *α*_mix_:

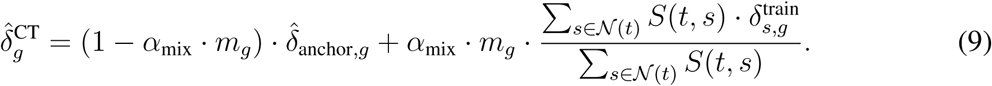

Genes outside the regulon (*m_g_* = 0) retain the structure-anchored prediction unchanged.

#### Nearest-neighbor supplement

Because cross-TF transfer operates only on the local regulon (genes with scaffold evidence), we introduced a complementary transcriptome-wide correction. Whereas the cross-TF transfer is mechanism-driven and localized to physically supported target genes, the nearest-neighbor supplement provides a global phenotypic anchor across all *G* genes. We retrieved the single training TF *s*^∗^ whose observed response profile is most correlated with the structure-anchored prediction in the full gene space:

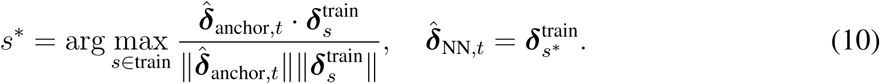

#### End-to-end prediction module

The final output blends the local mechanism-driven correction with the global phenotypic anchor:

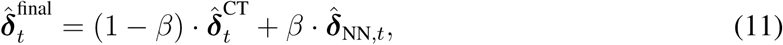

where *β* = 0.3 controls the weight of the transcriptome-wide supplement.

### Datasets

These four datasets collectively evaluate three distinct dimensions of the VitaGRN framework: (1) GRN topology quality and zero-shot perturbation response prediction on the K562 main benchmark, (2) cross-cell-line transferability via K562→RPE1 zero-shot transfer of both the regulatory scaffold and the response predictor, and (3) cross-task boundary characterization via GDSC2 drug response and sci-Plex chemical perturbation prediction. The combination of these datasets tests whether a physically grounded GRN retains its value beyond the cell line and perturbation class on which it was constructed.

#### K562 main benchmark dataset

The K562 dataset was obtained through the CausalBench preprocessing framework^33^ built on the Replogle et al. Perturb-seq system^2^. After our downstream filtering and feature selection, the working matrix comprised 152,060 perturbed single cells, 10,691 control cells, 1,092 perturbation labels, and 5,000 highly variable genes. For each perturbation *p*, the response vector was defined as the perturbation-level mean expression minus the control mean expression:

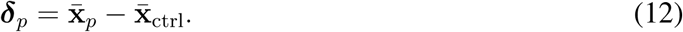

Control and perturbed expression profiles were stored as separate expression matrices for scaffold construction and response prediction. The scaffold construction pipeline additionally used a pre-computed baseline expression table for TF expression filtering and a cell-type-specific control expression profile for co-expression computation. Protein structure embeddings for 1,087 TFs were generated using the pre-trained ProTrek model^23^ as 1,024-dimensional structurally aligned vectors. TF–TF structural similarity edges were pre-computed as cosine similarities of these embeddings. The protein–protein interaction network was derived from STRING v12^32^ (combined score ≥ 400).

#### RPE1 cross-cell-line transfer dataset

The RPE1 dataset was obtained via the scPerturb harmonized resource^34^ from the same Perturb-seq experimental system^2^. After our filtering and feature selection, the working RPE1 matrix comprised 247,914 single cells, 2,393 perturbations, and 8,749 genes. This dataset was used exclusively to test whether the regulatory scaffold constructed in K562 and the response predictor could transfer to another cell line without retraining.

#### H9 developmental cell-state transition dataset

To evaluate VitaGRN’s capacity to capture dynamic temporal GRN rewiring along a continuous developmental trajectory, we utilized transcriptomic and genomic tracks from H9 human embryonic stem cells (ESCs) and H9-derived neural progenitor cells (NPCs). Matched steady-state RNA-seq data were obtained from an H9 ESC-to-NPC differentiation dataset generated using a dual-SMAD neural induction protocol, in which PAX6 marks the neuroectoderm transition^29^. For each cell state, the regulatory scaffold was constructed independently using sequence-level mutagenic scoring (ISM) and chromatin accessibility profiles. The dynamic rewiring of transcription factor (TF) targets between ESC and NPC states was quantified using the Jaccard distance of their target gene sets. For each TF, target sets *T*_ESC_ and *T*_NPC_ were defined by thresholding context-specific edge weights at *τ* = 0.01. The Jaccard distance is calculated as:

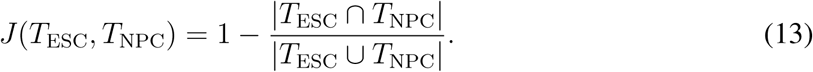

TFs were classified into four regulatory modes based on their expression change (log_2_ fold-change of NPC vs. ESC) and Jaccard distance, using thresholds *θ*_FC_ and *θ_J_* (see Supplementary Table 6): Master Drivers (| log_2_ FC| ≥ *θ*_FC_, *J* ≥ *θ_J_*), Chromatin Rewirers (| log_2_ FC| *< θ*_FC_, *J* ≥ *θ_J_*), Expression Followers (| log_2_ FC| ≥ *θ*_FC_, *J < θ_J_*), and Stable Regulators (| log_2_ FC| *< θ*_FC_, *J < θ_J_*).

To prioritize candidates for experimental validation, TFs were ranked by a Composite Prioritization Score (*S*_comp_) that aggregates target rewiring magnitude, edge confidence, state expression specificity, and biological novelty. Specifically, the score for TF *i* is defined as the geometric mean of its normalized criteria ranks:

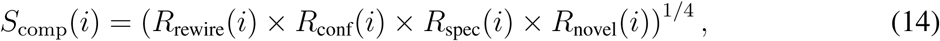

where *R*_rewire_(*i*) is the percentile rank of Jaccard target rewiring distance, *R*_conf_(*i*) is the percentile rank of mean scaffold edge confidence (*ϕ*^ˉ^_dom_), *R*_spec_(*i*) is the percentile rank of expression change magnitude (| log_2_ FC|), and *R*_novel_(*i*) is the percentile rank of biological novelty (prioritizing under-studied factors and non-canonical regulators lacked classical DNA-binding domains).

### Experimental details

#### Evaluation metrics

We report macro-averaged AUPRC as the primary ranking metric for GRN topology, alongside AUROC, recall, EP@K early-precision metrics, and edge direction sign consistency. For perturbation response prediction, we adopted DEG-focused metrics as the primary evaluation criterion (top-20 and top-50 PCC and direction consistency) because CRISPRi perturbations typically affect only a small number of highly responsive genes; genome-wide metrics (PCC-*δ*, Spearman, MAE, RMSE) are also reported.

#### Fold-safe split and perturbation benchmarking

The main response prediction evaluation adopted a perturbation-level fold-safe split for both K562 and RPE1 (consisting of train, validation, and test partitions), reserving 218 TF perturbations as the strictly held-out test set for K562. These TFs remained completely unseen during all training and hyperparameter tuning stages. To benchmark prediction performance, VitaGRN was compared against Matching Mean, GEARS, biolord, and a Latent Additive model under a unified protocol: all model outputs were aggregated to perturbation-level mean expression on a fixed 5,000-gene universe, and evaluated on delta vectors against the control mean. The fixed random seed and search space are provided in Supplementary Table 6. Regulatory scaffold topology evaluation was conducted within a unified matched candidate space, using a four-tier hierarchical gold standard: Tier 1 (CollecTRI) provides manually curated functional regulatory relationships; Tier 2 (BroadGRN) superimposes TRANSFAC motif filtering; Tier 3 (functional_all) aggregates CollecTRI, TRRUST^35^, and BroadGRN; and Tier 4 (+ ENCODE K562 ChIP-seq) adds cell-line-specific binding evidence used only for K562 sensitivity analysis. RPE1 evaluation used Tiers 1–3, because Tier 4 contains K562-specific binding evidence. The gold standard sources include CollecTRI^36^, TRANSFAC^37^, ENCODE K562 ChIP-seq, STRING v12^32^, and HOCOMOCO v11^31^; tier sizes, TF counts, and edge counts are provided in Supplementary Tables 7 and 8.

#### GRN topology evaluation framework

GRN topology evaluation was conducted within a fixed-space benchmarking framework on the K562 cell line. To ensure a fair comparison, the evaluation universe was restricted to the main candidate directed edge space (S_main_) and evaluated against the integrated physical-binding reference standard (G_ref_). We compared VitaGRN against a cohort of 20 other GRN inference models. Detailed benchmarking rules, including candidate space partitions, calibration rules, configuration audits, and auxiliary gold standard layers, are provided in the Supplementary Information.

### Statistics and reproducibility

#### Statistical significance and reproducibility

Pairwise statistical significance for GRN topology was determined using one-sided Wilcoxon signed-rank tests across TF-level AUPRC values with Benjamini–Hochberg correction. For direction accuracy and other sample-level estimates, 95% confidence intervals were derived using bootstrap resampling (1,000 resamples). Random-topology controls and dataset-dropout robustness were conducted across 5 random seeds to verify stability.

#### Computational resources and execution time

All experiments were conducted on a single server equipped with NVIDIA H20 GPUs. Regulatory scaffold construction (offline AlphaGenome ISM prediction) is a one-time computation requiring several GPU-hours; model fitting and virtual perturbation inference are both highly efficient.

### Data availability

All datasets used in this study are publicly available. K562 Perturb-seq data were obtained from the CausalBench repository (https://github.com/causalbench/causalbench). RPE1 Perturb-seq data and sci-Plex data were obtained from the scPerturb standardized resource (http://scperturb.org). GDSC2 drug response data were obtained from the DepMap Portal (https://depmap.org/portal/). Gold standard regulatory relationships were sourced from Collec-TRI (https://github.com/saezlab/CollecTRI), TRRUST (https://www.grnpedia.org/trrust/), BroadGRN (https://maayanlab.cloud/Harmonizome/), DoRothEA (https://saezlab.github.io/dorothea/), ENCODE K562 ChIP-seq (https://www.encodeproject.org/), and STRING v12 (https://string-db.org/). TF binding motifs were sourced from HOCOMOCO v11 (https://hocomoco11.autosome.org/) and JASPAR CORE 2024 (https://jaspar.elixir.no/). The following datasets gen-erated in this study have been deposited on HuggingFace at https://huggingface.co/ `datasets/Chris-young-2004/VitaGRN` and are publicly available: (1) The K562 *In Silico* Perturbation Atlas (containing the predicted Δ matrix and Bayesian uncertainty *σ* matrix for 1,087 TFs over 5,000 genes); (2) The VitaGRN biophysical regulatory scaffold (246,584 directed edges); (3) The structurally aligned ProTrek embeddings specifically generated for the 1,087 evaluated TFs; (4) Evaluation gold standard libraries (Tiers 1–4) and train/validation/test splits used for the benchmarking framework. Source data for all main-text and extended data figures are provided with the manuscript.

### Code availability

All original code, including the VitaGRN inference framework, the trained BayesianRidge model weights, and the evaluation scripts, has been open-sourced on GitHub at https://github.com/zwYang2004/VitaGRN and is publicly available.

## Acknowledgements

We thank all members of the AI Laboratory at Westlake University for helpful comments and discussions.

## Author contributions

Z.Y. and S.Z.L. conceived the study. Z.Y. designed the core algorithmic framework of VitaGRN, performed the formal analyses and wrote the manuscript. S.H. and G.B. performed the comprehensive benchmark evaluations and contributed equally. J.D. contributed to early-stage framework validation and methodological exploration. J.W. contributed to paper revision and analysis. S.Z.L. supervised the research, provided resources and acquired funding. All authors read and approved the final manuscript.

## Additional information

**Supplementary Information** is available for this paper.

## Extended Data Figures

The following 8 Extended Data Figures supplement the main text analyses. All data come from the unified evaluation pipeline; values are consistent with the main text.

**Extended Data Fig. 1.**
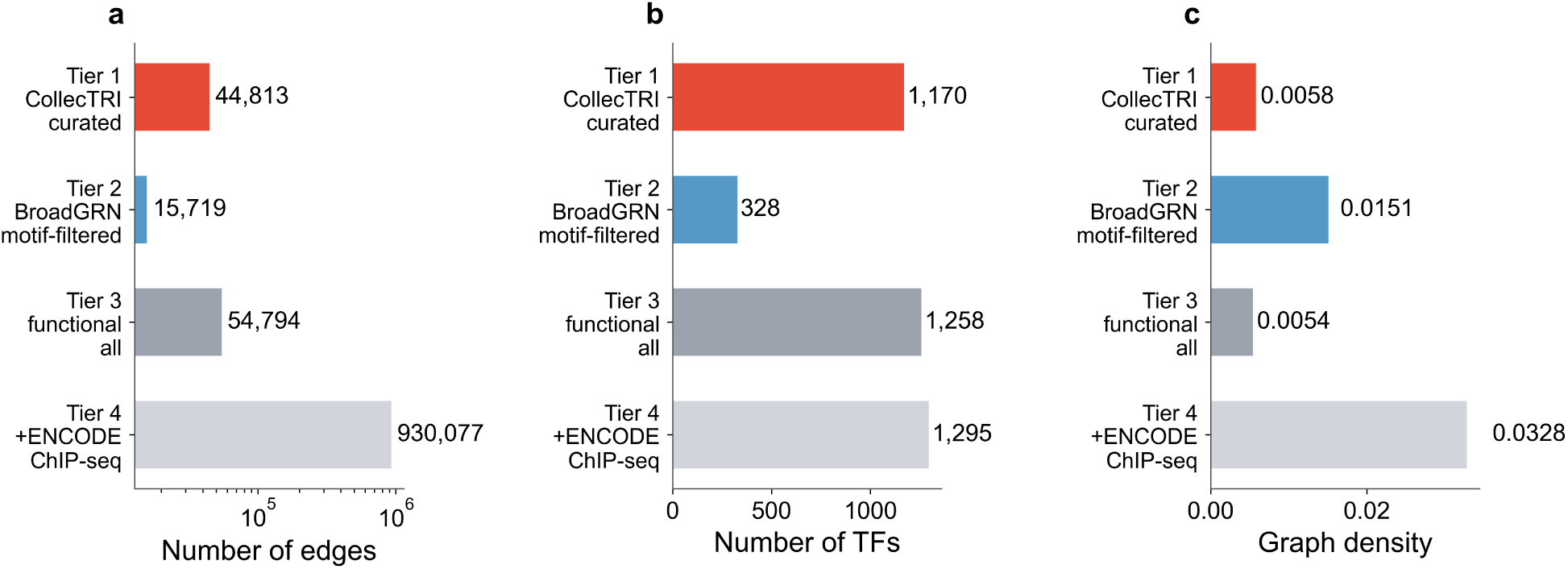
| Four-tier Gold Standard Library. **a**, Edge counts (log scale) across the four gold standard tiers. **b,** Number of transcription factors (TFs) represented in each tier. **c,** Graph density for each tier. Tier 2 (BroadGRN) has the lowest density (0.0150, or 0.01% rounded in text) among typical functional reference databases, providing the highest AUPRC discrimination.

**Extended Data Fig. 2.**
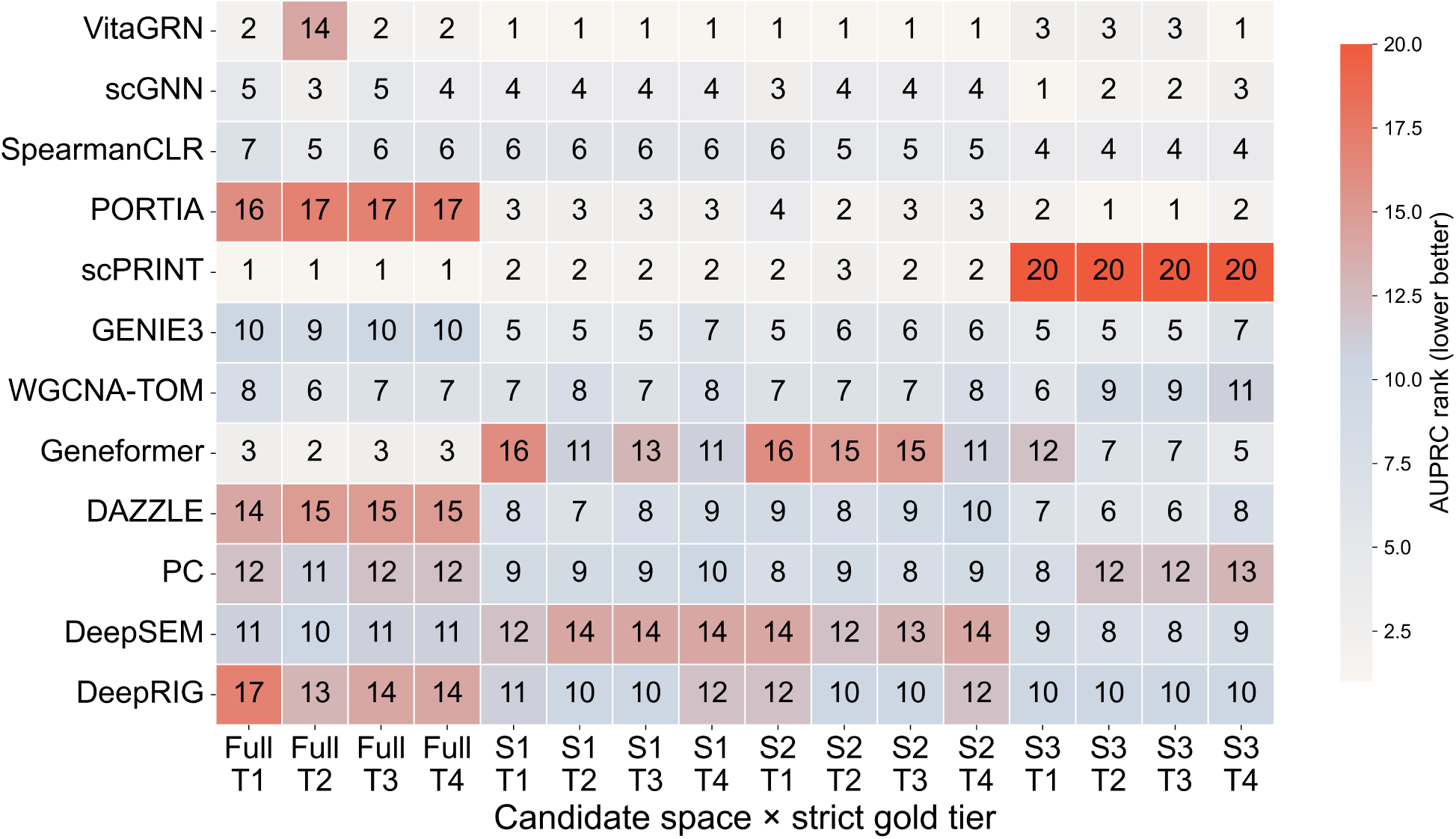
| Supplementary rank heatmap across K562 strict blocks. The heatmap reports model rank structures in non-primary strict candidate spaces and gold standard intersections for tie and boundary auditing, serving to define the sensitivity boundaries of the benchmark.

**Extended Data Fig. 3.**
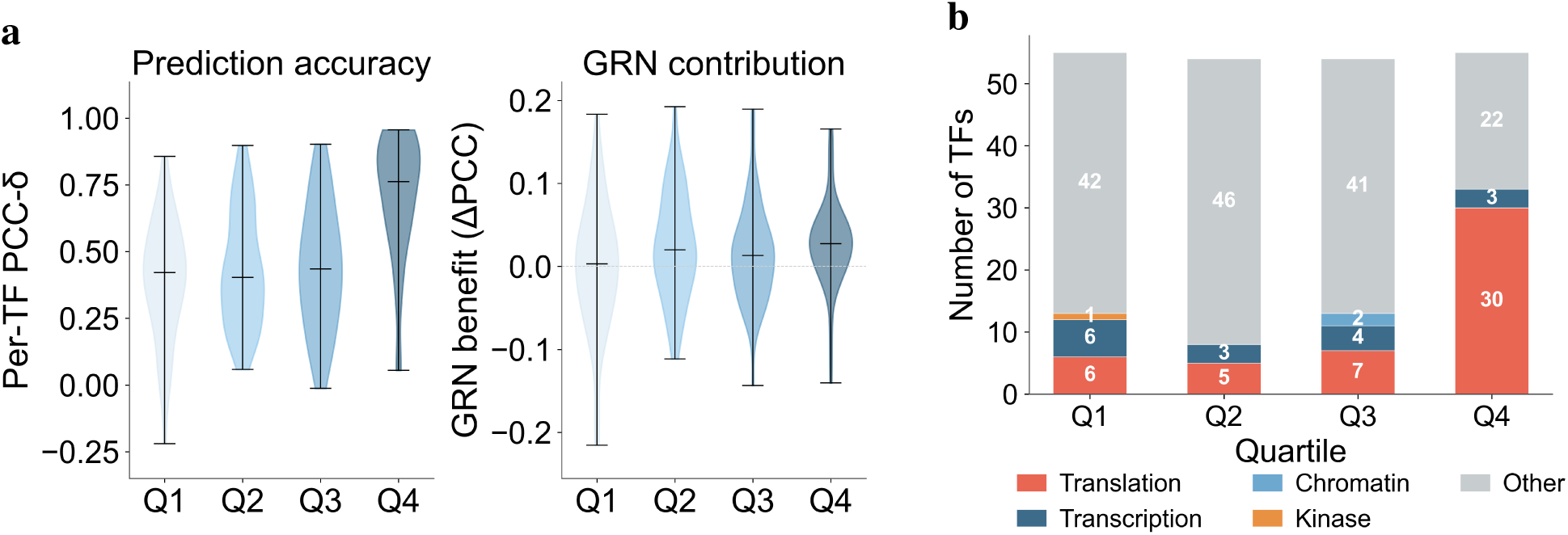
| Prediction quality correlates with perturbation effect size and functional category. **a**, Violin plots demonstrating that both prediction accuracy (left) and GRN contribution ΔPCC (right) increase significantly from the lowest-effect quartile (Q1) to the highest-effect quartile (Q4) of perturbation effect size. **b,** Transcription factor functional category distribution across quartiles: translation factors dominate the highly predictable Q4 group, while diverse signaling factors (“Other”) dominate Q1–Q3.

**Extended Data Fig. 4.**
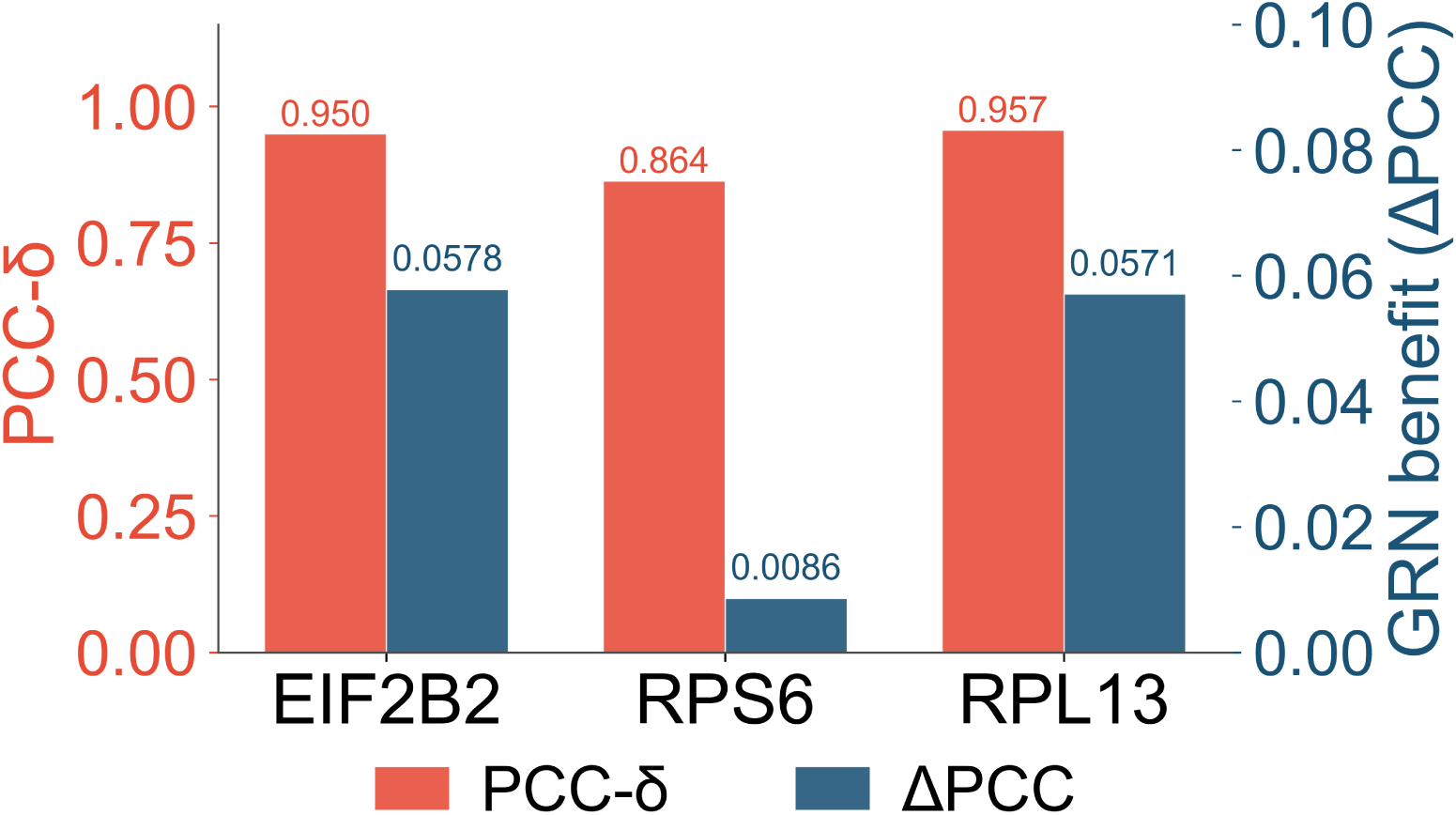
| Translation factors exhibit exceptional prediction accuracy and GRN contribution. Bar chart comparing prediction accuracy (PCC-*δ*, red) and GRN contribution (ΔPCC, blue) for translation and ribosomal factors (EIF2B2, RPS6, RPL13). The physical regulatory network injects substantial performance gains (e.g., +0.058 for EIF2B2) into these non-canonical targets.

**Extended Data Fig. 5.**
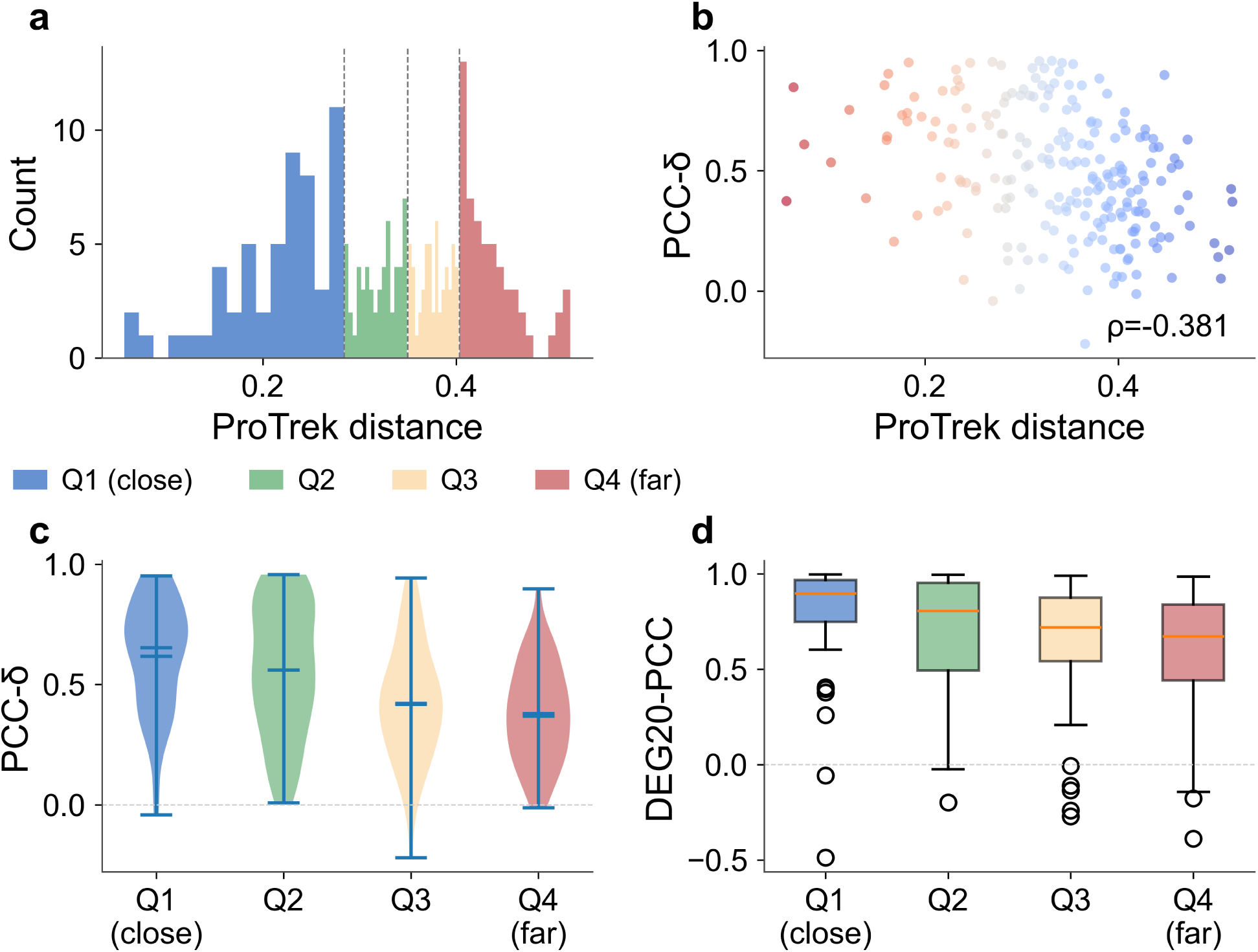
| Zero-shot Split Strictness. **a**, ProTrek cosine distance distribution to the nearest training TF for the 218 test TFs (quartile-colored). **b,** Scatter plot showing prediction accuracy (PCC-*δ*) against ProTrek structural distance. **c,** Violin plots of per-TF prediction accuracy (PCC-*δ*) across structural distance quartiles. **d,** Box plots of per-TF generalization accuracy (DEG20-PCC) across structural distance quartiles. Q4 remains significantly above random, proving true generalization.

**Extended Data Fig. 6.**
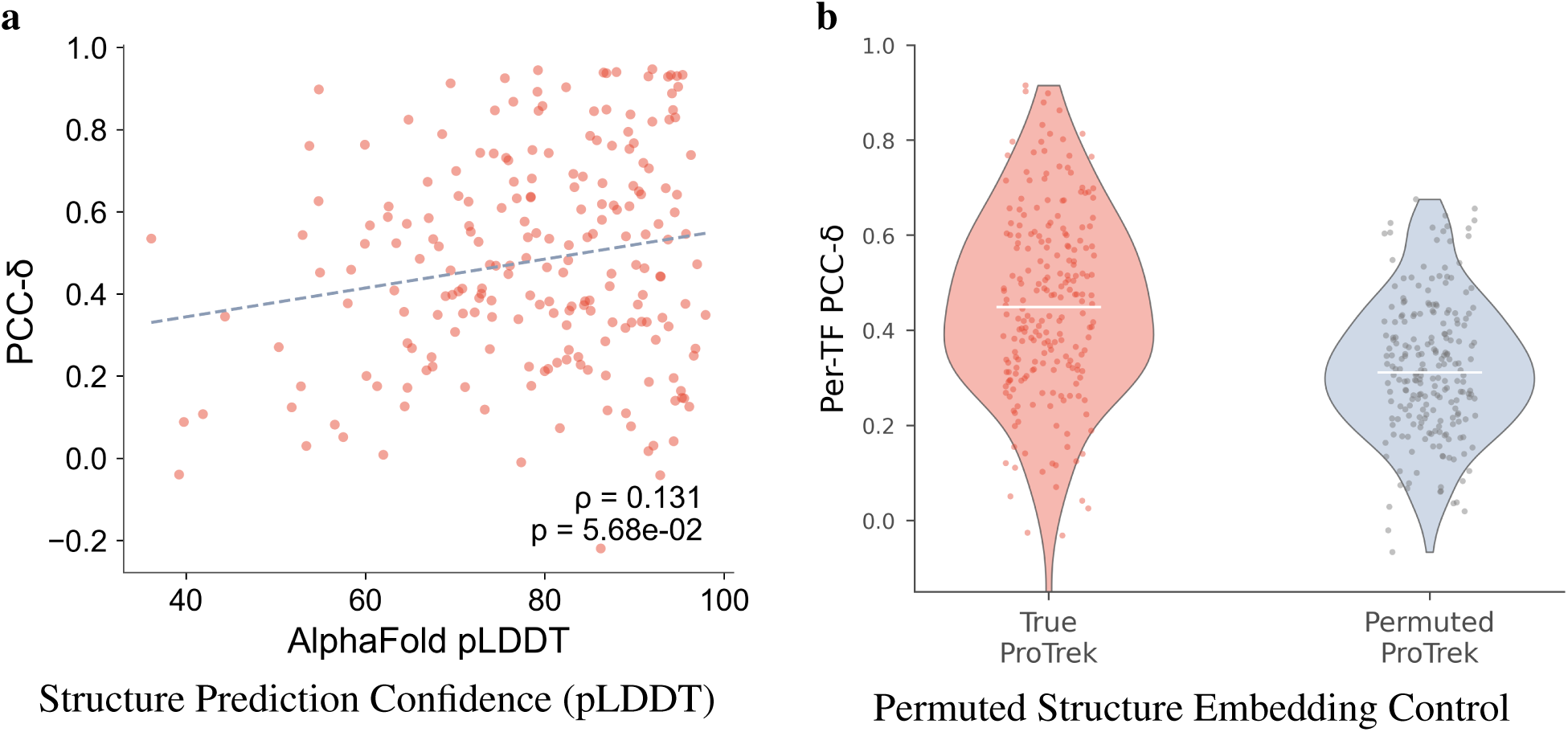
| Protein structure confidence and embedding validation. **a**, Scatter plot demonstrating that prediction accuracy (PCC-*δ*) is independent of AlphaFold structural prediction confidence (pLDDT). **b,** Negative control violin plots comparing prediction accuracy under true versus permuted ProTrek structure embeddings, showing a collapse in prediction quality when the TF-to-structure correspondence is broken.

**Extended Data Fig. 7.**
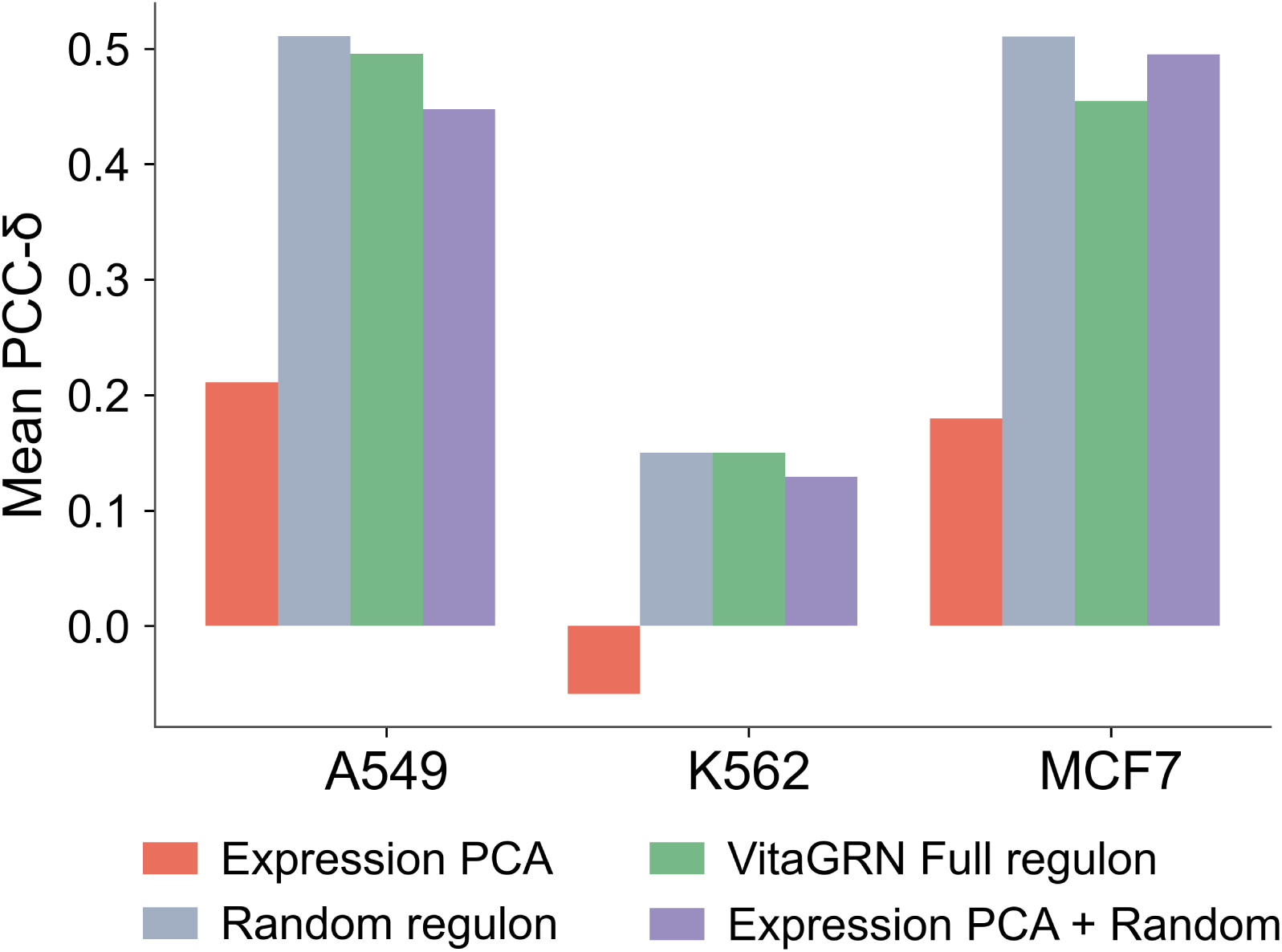
| Downstream Task Transfer Boundary Full Analysis. Comparison of prediction accuracy (Mean PCC-*δ*) across three cell lines (A549, K562, MCF7) under four feature configurations (Expression PCA, Random regulon, VitaGRN Full regulon, and Expression PCA + Random) from the sci-Plex perturbation dataset. This analysis explores the model’s performance boundaries and transfer limitations under complex chemical perturbations.

**Extended Data Fig. 8.**
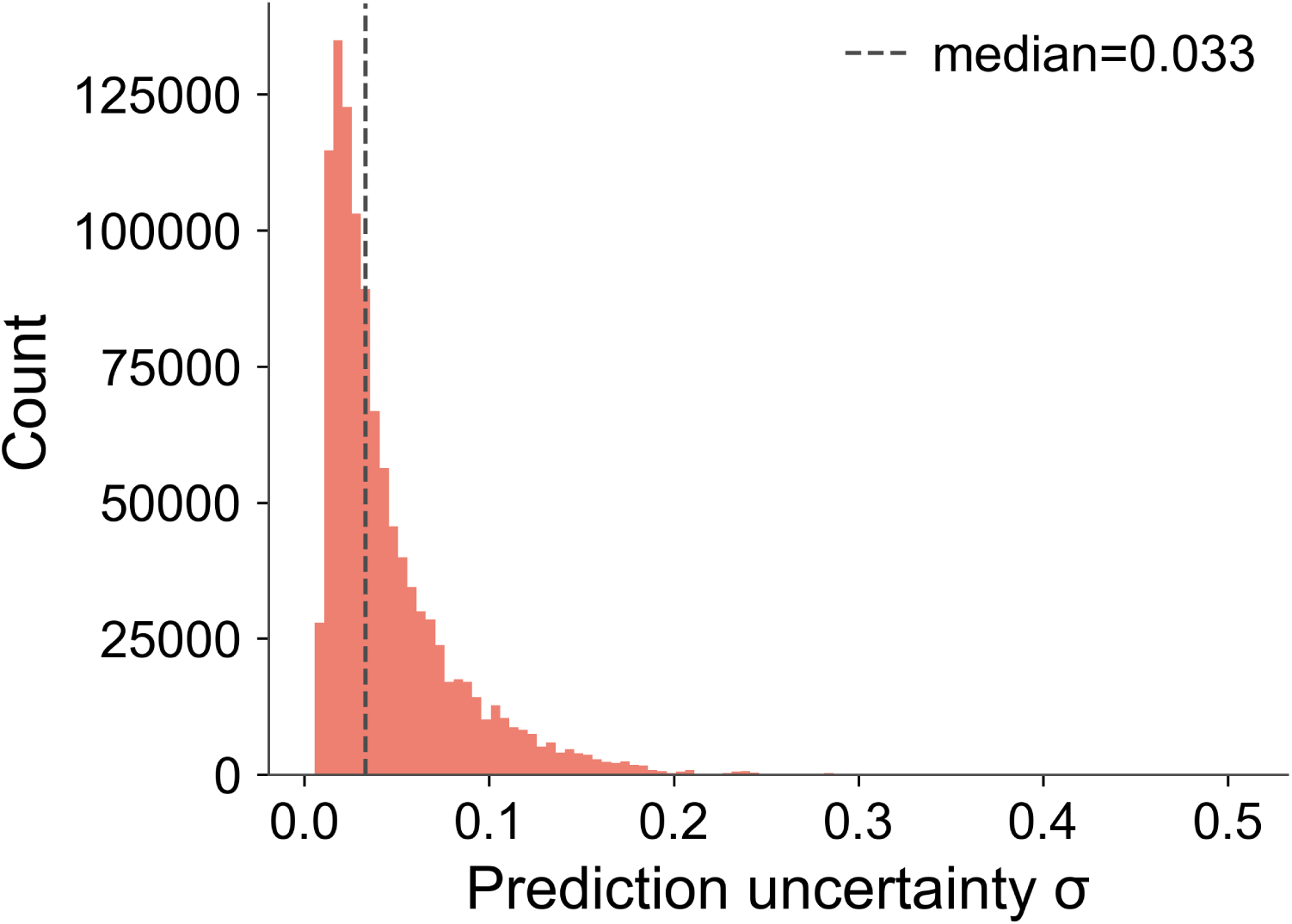
| BayesianRidge uncertainty quantification. Histogram showing the distribution of uncertainty (*σ*) estimates across all transcription factor–target gene pairs in the K562 perturbation database. The dashed line indicates the median uncertainty.

