## Supplementary Information for "Multimodal physical evidence uncovers interpretable gene regulatory networks for perturbation prediction"

### Supplementary Methods

#### Supplementary Method 1: K562 Fixed-Space Gene Regulatory Network Benchmarking Framework

**Overview of the fixed-space benchmark.** We evaluated VitaGRN in a K562 GRN benchmark that separates model prediction from evaluation-space definition. The benchmark first fixes the input universe, candidate-space partition, gold-standard hierarchy, and scoring rules, and then compares all methods within the same directed TF–target edge schema. This design is intended to answer a specific question: given the same K562 candidate universe and the same regulatory reference, which method assigns higher ranks to validated regulatory edges?

The main-text evaluation is performed on the main candidate space ($\mathcal{S}_{\text{main}}$) against the integrated reference standard ($\mathcal{G}_{\text{ref}}$). Here, $\mathcal{S}_{\text{main}}$ fixes the denominator of candidate TF–target edges, and $\mathcal{G}_{\text{ref}}$ fixes the positive reference set. Other candidate spaces and reference standard definitions are retained for sensitivity analysis, but they are not used to redefine the primary claim.

**Evaluation workflow.** We formulated the K562 gene regulatory network (GRN) benchmark as a directed TF–target edge-ranking problem within a fixed candidate universe. For each evaluated method, the input is a scored set of candidate regulatory edges; the objective is not to redefine the evaluable boundary, but to rank evidence-supported regulatory edges ahead of the remaining candidates under the same K562 denominator. Before scoring, we prespecified the candidate space, the gold-GRN evidence layer, the edge-canonicalization rules, and the reporting hierarchy. All predictions were then projected onto a common directed-edge schema and evaluated under the same space–gold block. This design follows the central benchmarking principle that GRN methods should be compared on a shared background universe and a shared positive reference set rather than on method-specific denominators[1].

**Candidate spaces and primary-space selection.** The K562 benchmark is not a single expression matrix, but an evaluation object jointly defined by ranked prediction lists, candidate TF–target universes, and stratified gold GRNs. For a candidate space $\mathcal{S}$ and gold GRN $\mathcal{G}$, positives are defined as $\mathcal{S} \cap \mathcal{G}$; all other candidate edges remain unlabeled background under the current evidence layer rather than confirmed biological false positives.

We retained three K562 candidate spaces to separate coverage, comparability, and intersection effects:

- **Main candidate space ($\mathcal{S}_{\text{main}}$)**: Formulated as the union of predicted edges across the VitaGRN model family. This primary space contains 5,396 genes, 1,078 candidate TFs, and 5,816,888 candidate directed edges. It balances gene/TF coverage and evaluable positive density, and is used as the primary candidate universe for all main evaluations. It preserves the submitted VitaGRN search domain while removing the unconstrained TF background of the full universe.

- **Shared candidate space ($\mathcal{S}_{\text{shared}}$)**: Formulated as the intersection of predicted edges across models within the family. It contains 3,293 genes, 978 candidate TFs, and 3,220,554 candidate directed edges, serving as a robustness analysis space. It imposes a tighter shared denominator across the VitaGRN family while retaining enough projected positives for stable evaluation.

- **Global candidate space ($\mathcal{S}_{\text{global}}$)**: The unconstrained genomic background containing 5,968 genes, 5,966 candidate TFs, and 35,605,088 candidate directed edges before gold projection. It preserves the largest background and is used as a denominator control.

Selecting $\mathcal{S}_{\text{main}}$ fixes the primary evaluation denominator, ensuring that newly submitted or externally generated models are scored on a unified benchmark universe. Details of space sizes and projected gold coverage are reported in Supplementary Tables 8 and 14.

**Prediction cohort and canonical edge representation.** The evaluation cohort contains 21 GRN prediction models, including submitted models and external comparators. The comparator panel represents diverse GRN-inference ideas, including tree-based inference, mutual-information or correlation networks, co-expression networks, regulon inference, perturbation or network-simulation approaches, and graph/deep-learning models[2–10]. To make heterogeneous outputs comparable, each prediction file was canonicalized as

$$e = (\mathrm{TF}, \mathrm{Target}, s_e),$$

where $s_e$ is the ranking score. Gene symbols were harmonized to HGNC-approved nomenclature before scoring[11]. Self-loops were removed, and duplicated TF–target pairs were collapsed to a single candidate edge. Thus, within any evaluation block, each method contributes at most one ranking score for each evaluable directed edge.

**Comparator-configuration fairness and score calibration.** Comparator fairness was enforced at two levels. First, the prediction cohort was scored through a single evaluator: all models were projected to the same main candidate space $\mathcal{S}_{\text{main}}$, intersected with the same integrated reference standard $\mathcal{G}_{\text{ref}}$, and evaluated by the same metric code path. Second, model-generation metadata were audited before interpretation. The available audit separates models evaluated from fixed released edge lists from models rerun with recorded metadata, and from models whose threshold or retained-edge budget was selected on validation folds. For rerun models with validation-based selection, the selection rule used validation correlation rather than the test gold labels, and the selected configuration was then frozen before $\mathcal{G}_{\text{ref}}$ scoring. We therefore describe the benchmark as a fixed-space, fixed-evaluator comparison with audited configuration provenance, rather than as a claim that every external method was exhaustively optimized over all possible hyperparameters.

Score calibration was handled by using ranking-based comparisons rather than cross-model score magnitudes. The unified loader treats larger regulatory magnitude as stronger evidence: signed weights are mapped to $s_e = |w_e|$ for ranking, while the original sign is retained as a separate signed score for sign-consistency auditing. Methods whose outputs are already non-negative confidence scores are ranked directly after this canonicalization. Because AUPRC, AUROC, and EP@$K$ depend on within-model ordering over the same TF-specific candidate targets, monotone transformations of a model's scores do not change its evaluated ranking. The benchmark therefore avoids comparing raw score scales across algorithms.

**Gold-GRN construction.** Gold GRNs were organized by evidence strata rather than pooling regulatory evidence with different reliability, provenance, and coverage into a single undifferentiated reference. The primary integrated reference standard ($\mathcal{G}_{\text{ref}}$) corresponds to the K562 Tier-4 all-source physical-binding layer (denoted $\mathcal{G}_{\text{ref}}$ in the pipeline). This layer integrates curated functional interactions (CollecTRI, TRRUST, BroadGRN) with cell-line-specific binding evidence (ENCODE ChIP-seq), treating physically supported TF–target edges as the positive reference. It was selected because it preserves direct binding support while retaining sufficient positive-edge density for stable ranking.

Auxiliary gold definitions were used only for sensitivity analysis and were not allowed to redefine the primary endpoint. These include single-source references (CollecTRI, TRRUST, BroadGRN, and ENCODE K562 ChIP-seq), multi-source intersection layers, and a high-confidence expanded layer incorporating DoRothEA A–C evidence[12]. The high-confidence expanded layer admits an edge if it is supported by at least two functional sources, or if one functional source is accompanied by ENCODE K562 binding evidence. All gold layers are intersected with the candidate space before scoring; therefore, raw evidence-library size and the projected scoring size are reported separately (Supplementary Table 7).

**Primary task, metrics, and reporting hierarchy.** The primary evaluation task measures model performance in the main candidate space ($\mathcal{S}_{\text{main}}$) against the integrated reference standard ($\mathcal{G}_{\text{ref}}$). All main-text comparative claims are restricted to this configuration. Analyses using the global space ($\mathcal{S}_{\text{global}}$), the shared space ($\mathcal{S}_{\text{shared}}$), or auxiliary reference layers are reported as robustness controls in the appendix.

The primary ranking criterion is macro-averaged AUPRC. For each evaluated gold TF, predictions are ranked over its candidate targets and a TF-specific precision–recall curve is computed; these values are then averaged across gold TFs:

$$\mathrm{AUPRC}_{\mathrm{macro}} = \frac{1}{|\mathcal{F}_{\text{gold}}|} \sum_{f \in \mathcal{F}_{\text{gold}}} \mathrm{AUPRC}_f.$$

This macro formulation prevents TFs with many gold edges from dominating the benchmark. Secondary metrics include AUROC, AUPRC-lift, recall, and early-budget precision:

$$\mathrm{AUPRC}_{\mathrm{lift}} = \frac{\mathrm{AUPRC}}{\rho_{\mathrm{gold}}}, \qquad \mathrm{EP@}K = \frac{\mathrm{TP@}K}{\min(K, N_{\mathrm{pred}})}.$$

For sparse or very small blocks, we use minimum-rank tie assignment and retain complete tie and coverage audits in the appendix.

**Uncertainty estimation and paired significance testing.** Uncertainty was quantified only for the prespecified primary block. Because the primary metric is macro-averaged over gold TFs, resampling was performed at the TF level: gold TFs were sampled with replacement and macro-AUPRC was recomputed for each bootstrap replicate. Percentile intervals from 1,000 bootstrap replicates were used as 95% confidence intervals for each model. Pairwise comparisons between VitaGRN and each comparator used paired TF-level AUPRC values in a one-sided Wilcoxon signed-rank test, with Benjamini–Hongberg correction across comparator tests. These analyses quantify uncertainty in the same unit used by the macro endpoint and do not alter the prespecified primary task.

#### Supplementary Method 2: Independent perturbation validation protocols

The main text reports core metrics for six perturbation data validation tasks. The following sections provide supplementary protocol details.

**Target gene recovery protocol.** For each held-out TF, its observed top-50 DEGs (ranked by $|\delta|$) serve as positives; the remaining genes among the 5,000 are negatives. Ranking scores use absolute GRN edge weights (for regulatory scaffold methods) or absolute predicted delta (for end-to-end predictions). AUROC and AUPRC are averaged across 218 TFs.

**Fair direction accuracy protocol.** To avoid evaluation bias from differing numbers of evaluable TFs across methods, we define each method's evaluable TF set as those simultaneously satisfying (i) $\geq$10 signed edges overlapping DEGs and (ii) at least one positive and one negative edge. Direction accuracy is recomputed on the intersection of evaluable TFs shared by at least two methods, using bootstrap 95% CI (1,000 resamples).

**Effect size correlation protocol.** PCC-$\delta$ is computed on genome-wide perturbation-level delta (5,000 genes). First-order propagation baselines predict the response of candidate target gene $g$ to TF $x$ knockdown as $-\mathrm{weight}(x \to g)$, with zero for genes without direct edges. End-to-end predictions use predicted delta directly.

**Novel regulatory relationship discovery protocol.** For each test TF, GRN edges not present in Tier 3 gold (54,794 edges) are designated "novel." Their target genes are checked against the TF's observed top-50 DEGs. The novel-to-gold validation rate ratio measures the perturbation support for newly discovered regulatory relationships relative to known ones.

**Pathway enrichment protocol.** For each TF's top-100 predicted targets, hypergeometric tests are performed against Reactome (752 gene sets), GO Biological Process (1,347), and KEGG (242), totaling 2,341 pathway gene sets, with BH-FDR correction (FDR $< 0.05$). The mean number of significant pathways per TF and the mean $-\log_{10}(p)$ of the most significant pathway are reported.

**Perturbation response retrieval protocol.** Given a query TF $q$ with observed perturbation response vector $\boldsymbol{\delta}_q^{\mathrm{obs}}$, the Pearson correlation between $\boldsymbol{\delta}_q^{\mathrm{obs}}$ and each candidate TF $c$'s predicted response $\hat{\boldsymbol{\delta}}_c$ is used as similarity to compute MRR, Hit@10, and pairwise AUROC (discriminating the positive pair $(q, q)$ from negative pairs $(q, c), c \neq q$). DEG-20 AUROC computes the discrimination AUROC using only each query TF's top-20 observed DEGs.

**Direction consistency evaluation protocol.**

Direction consistency measures the sign-match ratio between predicted regulatory edges and signed gold-standard annotations from CollecTRI[13] and DoRothEA[12]. VitaGRN edge weights are derived from perturbation direction or expression change direction and are therefore directly comparable with these signed references. For methods whose outputs consist of unsigned edge strengths, absolute correlation values, or non-negative importance scores, direction consistency is not applicable, because interpreting non-negative scores as activation would systematically inflate the metric. To prevent this, the evaluation pipeline checks whether predicted edge weights contain any negative values; if all scores are non-negative, the model is classified as unsigned and direction consistency is reported as n/a. This protocol was applied uniformly across all methods in both the K562 and RPE1 evaluations.

### Supplementary Method 3: Detailed perturbation response benchmark protocol

**Benchmark scope.** This benchmark evaluates transcription factor (TF) perturbation response prediction on K562 cells under a unified generalization protocol where test TFs are completely held out. The goal is to compare VitaGRN against established external baselines under identical test perturbations and gene universes. The comparison includes VitaGRN, Matching Mean, GEARS, biolord, and a Latent Additive model.

**Data source and fixed split.** The benchmark uses K562 Perturb-seq data obtained from essential-scale CRISPRi screens. The dataset is partitioned using a fixed split where 218 TFs are held out as a test set. These test perturbations are excluded from all model training, hyperparameter tuning, and post-hoc calibration steps to ensure unbiased evaluation of model generalization.

**Gene universe and control statistics.** All compared models are evaluated on a fixed 5,000-gene universe. Let $\bar{x}_{\mathrm{ctrl}} \in \mathbb{R}^G$ denote the control mean expression vector ($G = 5000$). For perturbation $p$, observed and predicted perturbation-level mean expression vectors are denoted by $\bar{x}_p$ and $\hat{x}_p$, respectively. Delta profiles are defined as

$$\Delta_p = \bar{x}_p - \bar{x}_{\mathrm{ctrl}}, \quad \hat{\Delta}_p = \hat{x}_p - \bar{x}_{\mathrm{ctrl}}.$$

**Unified prediction format.** To ensure a standardized comparison, predictions from all models are formatted into a unified structure, containing predicted expression values, observed ground-truth values, perturbation identities, and control baseline expression. For models that produce non-standard output representations, standard adapters are used to align their outputs to this common format without modifying the underlying model architectures, ensuring fair and consistent evaluation.

**Model reproduction and adaptation.** All models are evaluated at the perturbation level using the same evaluation pipeline across the fixed held-out split and the 5,000-gene universe, ensuring that no information from test perturbations is used during training or hyperparameter selection.

*VitaGRN.* The evaluation uses the predictions generated by VitaGRN under the weighted cosine hybrid configuration. The final predictions are evaluated under the same pipeline as the baseline methods, covering the complete set of 218 test perturbations ($n_{\text{eval}} = 218$).

*Matching Mean.* Matching Mean serves as a non-parametric baseline representing the average response profile across all training perturbations. Under the evaluation protocol, the observed training perturbation responses are averaged to form a reference profile, which is then used as the prediction for all held-out test perturbations. This baseline is evaluated under the same conditions ($n_{\text{eval}} = 218$) to assess the baseline predictive performance of shared, non-specific perturbation programs.

*GEARS.* GEARS is included as a graph-based baseline for perturbation response prediction. Due to gene ontology filtering during data preprocessing in GEARS, it evaluates 217 test perturbations ($n_{\text{eval}} = 217$), with the perturbation targeting C14ORF178 excluded from its evaluation.

*biolord.* biolord is included as a latent generative baseline. It is evaluated at the perturbation level on the same test set of 218 perturbations ($n_{\text{eval}} = 218$).

*Latent Additive.* The Latent Additive model is implemented as a baseline that predicts perturbation responses by adding perturbation vectors to control baselines in a low-dimensional latent space. Hyperparameters are selected strictly using the validation set, testing regularization strengths $\alpha \in \{0.01, 0.1, 1, 10, 100\}$ and selecting $\alpha = 100$. The model evaluates the full set of 218 test perturbations ($n_{\text{eval}} = 218$).

*Squidiff.* Squidiff was evaluated but excluded from the primary comparative tables due to a severe output-scale mismatch, where the predicted values do not align with the scale of the observed ground-truth expression, resulting in disproportionately high error metrics. In addition, the predicted outputs lack gene name identifiers, preventing unambiguous mapping of predictions to the target gene universe.

*Compositional Perturbation Autoencoder (CPA).* CPA was excluded from the zero-shot generalization benchmark because it relies on learning categorical perturbation embeddings for each target during training. Consequently, it is incapable of making predictions for completely held-out, unseen transcription factors.

**Perturbation-level aggregation.** Evaluation is perturbation-level. If a model outputs cell-level predictions, outputs are first aggregated to perturbation-level mean expression per perturbation, then transformed to delta against control mean. Models that directly output perturbation-level predictions are evaluated directly. This aggregation unifies output granularity across baseline families.

**Evaluation metrics.** Primary metrics are PCC-$\delta$, DEG20-PCC, DEG50-PCC, and DirMatch.

- PCC-$\delta$: for each perturbation $p$, Pearson correlation between $\hat{\Delta}_p$ and $\Delta_p$ over genes, followed by averaging over test perturbations.
- DEG20-PCC / DEG50-PCC: for each perturbation $p$, rank genes by $|\Delta_p|$ and select top-20/top-50; compute Pearson correlation between predicted and observed delta on that subset; average over perturbations.
- DirMatch: directional agreement between signs of predicted and observed delta profiles.

Supplementary metrics include LFCSpear and RMSE-$\delta$.

- LFCSpear: Spearman correlation between predicted and observed log-fold-change perturbation profiles.
- RMSE-$\delta$: root-mean-square error between $\hat{\Delta}_p$ and $\Delta_p$, aggregated across perturbations; lower is better.

Metric edge-case handling follows the standard implementation and is kept unchanged across all models.

**Benchmark reporting and exclusions.** The primary comparison includes VitaGRN, Matching Mean, GEARS, biolord, and Latent Additive. GEARS is evaluated on 217 perturbations due to gene ontology filtering, while other methods cover 218 perturbations. Squidiff and CPA are excluded from the main comparative results for the reasons detailed above.

**Reproducibility and provenance.** To ensure reproducibility, all evaluation scripts, processed predictions for all compared baselines, and parameter configurations are hosted in the companion repository. Key outputs include the compiled metric summaries, visualization scripts, and model reproduction code.

### Supplementary Results and Discussion

#### Supplementary Note 1: Per-experiment result breakdown and confidence calibration

The following decomposes VitaGRN's core results by experimental dimension, supplementing details that could not be expanded in the main text due to space constraints.

**Complete GRN topology evaluation rank heatmap.** Extended Data Fig. 2 presents the complete unified rank heatmap for all methods across candidate spaces and gold standard evidence layers, visually summarizing the global performance landscape under the K562 benchmark.

**Complete perturbation response prediction benchmark.** Supplementary Table 2 reports the quantitative comparison of VitaGRN against four baseline models (Matching Mean, GEARS, biolord, and Latent Additive) across six complementary evaluation metrics. VitaGRN outperforms all baseline methods across every metric: achieving a PCC-$\delta$ of 0.495 (compared to the second-best Latent Additive at 0.373 and Matching Mean at 0.370), a DEG20-PCC of 0.681 and DEG50-PCC of 0.686 (compared to Latent Additive at 0.557 and 0.570), a direction match (DirMatch) of 0.635, a log-fold change Spearman correlation (LFCSpear) of 0.370, and a reduced root-mean-squared error (RMSE-$\delta$) of 0.076. These results demonstrate that the integration of the biophysical scaffold prior consistently improves both correlation and direction accuracy in quantitative perturbation prediction.

**Response prediction per-TF distribution.** The main text reports mean metrics over 218 held-out TFs. The per-TF DEG20-PCC median is 0.773 (Structure Anchor standalone), with a left-skewed distribution: most TFs have high prediction quality, with a small number performing poorly (minimum $\sim$0.2). These low-quality TFs typically exhibit: (1) lack of structurally similar neighbors in the training set (structure distance $>$ 0.45), (2) very few GRN edges in the regulatory scaffold ($<$ 10), or (3) very low expression in K562 with weak perturbation response signal.

**Structure distance gradient per-TF details.** Supplementary Table 10 lists per-metric means and standard deviations for the four distance quartiles. Q4/Q1 retention ratios: PCC-$\delta$ 61.6%, DEG20-PCC 74.8%, DirAcc 86.5%. All quartiles remain significantly above the random baseline, confirming the model's generalization ability.

**Target recovery and effect size per-TF comparison.** The per-TF AUROC distribution (VitaGRN end-to-end vs. regulatory scaffold edge weights vs. PSGRN) is shown in Extended Data Fig. 1. VitaGRN end-to-end predictions achieve mean AUROC = 0.893 (median = 0.927) across 218 TFs; regulatory scaffold edge weights achieve mean AUROC = 0.522 (near random 0.5). The effect size signed Spearman $\rho$ per-TF distribution is similar: end-to-end median $\rho$ = 0.540, regulatory scaffold ISM median $\rho \sim 0$.

**Fair direction accuracy across methods.** Direction accuracy comparisons across GRN methods are complicated by differential TF coverage: most GRN methods have signed edges for only a small subset of TFs, while VitaGRN's end-to-end predictions cover all 218 test TFs. To ensure a fair comparison, we restricted evaluation to the common ground of 96 TFs where at least two methods produce evaluable signed predictions (each having $\geq$10 signed edges overlapping observed DEGs). On this fair subset, VitaGRN achieves direction accuracy of 0.837 (bootstrap 95% CI [0.807, 0.866]), substantially exceeding the next-best methods (PORTIA: 0.506, PSGRN: 0.480). The margin is narrower than the all-TF comparison (0.862 vs. 0.480) because some hard-to-predict TFs are excluded from the common ground, but VitaGRN's advantage remains highly significant with non-overlapping confidence intervals.

**Uncertainty quantification calibration.** Extended Data Fig. 8 demonstrates the calibration capability of BayesianRidge $\sigma$. Per-TF mean $\sigma$ has a negative Spearman correlation with DEG20-

PCC, demonstrating that the model's uncertainty estimates are well-calibrated: TFs assigned low $\sigma$ are indeed predicted more accurately. High-confidence genes (bottom-50% $\sigma$) achieve significantly higher DEG20-PCC than low-confidence genes (top-50% $\sigma$), validating $\sigma$ as an effective metric for wet-lab prioritization.

**K562 *In Silico* Perturbation Atlas statistics.** The atlas covers 1,087 perturbation factors $\times$ 5,000 genes = 5,435,000 $\Delta$ predictions, each with $\sigma$ uncertainty, totaling 10,870,000 values. Generation time: 34 seconds (single CPU, BR inference). Three quality tiers: Tier 1 (779 TFs, training-visible + regulatory scaffold), Tier 2 (194 TFs, zero-shot + regulatory scaffold), Tier 3 (114 TFs, structure-only, no GRN). The mean absolute per-TF predicted $\Delta$ is 0.028 (consistent with typical delta magnitudes in K562 perturbation data); median $\sigma = 0.031$. The atlas is released as CSV files with the paper; single-TF queries can be performed using the provided Jupyter Notebook.

**Prediction quality stratified by perturbation effect magnitude.** We stratified the 218 test TFs into quartiles by predicted absolute perturbation effect (mean $|\Delta|$ across all 5,000 genes) to examine whether VitaGRN's prediction accuracy varies with effect magnitude. The Spearman correlation between predicted effect size and per-TF PCC-$\delta$ is $\rho = 0.41$ ($p < 10^{-4}$), confirming that larger-effect TFs are predicted more accurately. The GRN benefit ($\Delta$PCC = full GRN $-$ no GRN) is also concentrated in the upper quartile (see Supplementary Table 11). Medium-effect TFs (Q2) show the lowest prediction accuracy and the smallest GRN benefit, likely because they are enriched for signalling regulators (e.g. CHEK1, CPSF3) whose perturbation effects are distributed through multi-step cascades, producing complex transcriptional signatures that are more challenging to predict than the coherent responses of large-effect ribosomal or translation factors. This stratification is provided as a reference for users to calibrate expectations when querying the virtual atlas for TFs of differing effect magnitude.

**Virtual screening report and individual profile card.** The virtual perturbation screening report ($\sigma$ vs. $|\Delta|$ scatter plot) and the EIF2B2 individual profile card are shown in the main text (Figure 5a and Figure 3a, respectively).

#### Supplementary Note 2: Complete K562 GRN topology and system ablation results

**Data efficiency analysis.** Supplementary Table 9 summarizes performance changes as the training perturbation fraction varies from 5% to 100%. BranchA denotes Structure Anchor using SVD Embedding Ridge alone; Structure Anchor+NN adds nearest-neighbor transfer; Structure Anchor+GRN+NN adds weighted-cosine Cross-TF transfer guided by the VitaGRN regulatory scaffold.

**Extended biological case studies.** Supplementary Table 3 presents the 10 local biological case studies selected from the 218 held-out TFs. Selection criteria combine genome-wide PCC, DEG-focused PCC, the fraction of top DEGs covered by the VitaGRN local regulatory network, top-20 DEG direction consistency, and the canonical biological process associated with the perturbation.

#### Supplementary Note 3: Prediction quality and perturbation effect size analysis

To understand the biological drivers of prediction accuracy, we investigated the relationship between a transcription factor's perturbation effect size (the mean absolute magnitude of expression change across the genome, $|\Delta|$) and the model's prediction accuracy (per-TF PCC-$\delta$). As shown in **Extended Data Fig. 3a**, there is a clear positive correlation (Spearman $\rho = 0.41$) between the predicted effect size and actual prediction quality, indicating that TFs triggering massive systemic responses are fundamentally more predictable.

When stratifying the 218 test TFs into quartiles by their actual perturbation effect size from Q1 (smallest effect) to Q4 (largest effect) (**Extended Data Fig. 3a**), both the prediction accuracy and the marginal performance gain provided by the physical GRN prior ($\Delta$PCC) rise significantly. This demonstrates that for crucial regulators with pronounced biological effects, the VitaGRN biophysical scaffold contributes the core predictive advantage. Furthermore, analyzing the functional categories of the TFs within these quartiles (**Extended Data Fig. 3b**) reveals that while Q1–Q3 are dominated by diverse signaling and complex cascade factors (“Other”), the large-effect Q4 quartile is overwhelmingly dominated by translation and ribosomal factors (e.g., accounting for 25 of 55 factors). This explains why the model predicts translation factors with such high accuracy (**Extended Data Fig. 4**): these universally conserved macro-complexes have highly stable protein structures, and their perturbation triggers massive, coherent cellular stress responses (such as the integrated stress response) that VitaGRN’s structurally anchored GRN captures exceptionally well.

### Supplementary Note 4: RPE1 cross-cell-line GRN topology transfer evaluation

**Evaluation setup.** External GRN inference methods typically require cell-line-specific expression data as input and, by design, cannot transfer a K562-trained GRN to RPE1. VitaGRN directly projects its K562-constructed biophysical regulatory scaffold (246,584 edges) to the RPE1 gene space, without any RPE1-specific training or parameter adjustment. Evaluation used the unified evaluation pipeline across three cell-line-agnostic gold tiers (Tier 1 CollecTRI, Tier 2 BroadGRN, Tier 3 functional-all) and three matching candidate spaces (global space $\mathcal{S}_{\text{global}}$, shared space $\mathcal{S}_{\text{shared}}$, and main space $\mathcal{S}_{\text{main}}$).

**Results.** Supplementary Table 12 summarizes the key results. VitaGRN, using zero-shot transfer of the K562 GRN to RPE1, achieved significant AUPRC across all three gold tiers and all three evaluation spaces (Tier 1 CollecTRI = 0.0739, Tier 2 BroadGRN = 0.0407, Tier 3 functional-all = 0.0557; Tier 2 AUROC = 0.711, Recall = 0.507).

**Discussion.** VitaGRN’s core contribution lies at the paradigm level: the core layers of its biophysical regulatory scaffold (genomic sequence ISM and protein structure PPI cascade) are inherently cell-line-invariant, making “build once, reuse everywhere” physically feasible. Existing GRN methods, in their framework, require retraining when changing cell lines because the entire input data distribution changes. In this sense, VitaGRN opens a track that existing methods cannot enter: cross-cell-line zero-shot GRN generalization.

**Relationship to cross-cell-line response transfer.** The K562→RPE1 response prediction transfer (DEG20-PCC = 0.360) reported in the main text is a downstream extension of the GRN topology transfer documented here. GRN topology transfer demonstrates that the biophysical regulatory scaffold preserves regulatory relationship quality across cell lines; response transfer further demonstrates that this GRN, combined with the structure-guided response predictor, can generate meaningful cross-cell-line perturbation response predictions. The relationship between the two is illustrated in Figure 4f (cross-cell-line transfer results).

#### Supplementary Note 5: Boundary controls and structural robustness analyses

**Random topology control.** A critical boundary control tested whether the GRN's benefit depends on genuine regulatory topology. When the real GRN was replaced by randomly shuffled edges, prediction performance degraded, confirming that topological noise harms the structure-anchored prediction (**Figure 2e**). This confirms that integrating genuine biophysical prior topology is crucial for accurate quantitative perturbation predictions.

**Protein structure embedding validity.** A critical question for any structure-based method is whether the model genuinely exploits protein structure information or merely fits to spurious correlations in the embedding space. To test this, we permuted the structurally aligned protein embeddings of test TFs, breaking the correspondence between TF identity and structure, and re-evaluated prediction accuracy (**Extended Data Fig. 6b**). Under true embeddings, per-TF PCC-$\delta$ is broadly distributed (a mean of 0.450), with a long tail of well-predicted cases. Under permuted embeddings, the distribution collapses toward lower values (a mean of 0.319) and becomes tightly concentrated, indicating that the model loses its ability to discriminate between well- and poorly predicted TFs. This 29

**Structure prediction confidence.** We further tested whether protein structure prediction confidence (pLDDT scores, obtained from AlphaFold-predicted structures[14]) limits VitaGRN's performance (**Extended Data Fig. 6a**). Across 213 test TFs with available pLDDT scores, the correlation between structure confidence and prediction accuracy is weak ($\rho = 0.131$, $p = 0.057$). TFs with pLDDT as low as 40–50 achieve DEG20-PCC comparable to those with pLDDT $> 90$. This robustness to structural uncertainty is practically important: many disease-relevant proteins, particularly those with intrinsically disordered regions, have lower pLDDT scores, yet VitaGRN can still use their predicted structures as effective anchors for regulatory inference.

**Sparsity and noise robustness boundaries.** To evaluate the robustness of VitaGRN under input data limitations, we characterized its performance under sequence-level noise and single-cell sparsity. The regulatory scaffold relies on AlphaGenome ISM sequence scores as its biophysical foundation. Injected sequence-score noise led to a smooth, graceful decline in relative GRN AUPRC rather than a sudden collapse, stabilizing near 0.59 at 1.0–2.0$\times\sigma$ (**Figure 4g**). Similarly, to evaluate stability under cell subsampling, we measured response prediction performance across varying cell-dropout rates. Both PCC-$\delta$ and DEG20-PCC decreased smoothly with increasing dropout percentage, confirming that VitaGRN's response prediction is highly stable to input sparsity (**Figure 4h**). These analyses establish that the structure-anchored predictor remains reliable under typical experimental sequencing noise and shallow cell coverage.

#### Supplementary Note 6: H9 ESC-to-NPC edge weight validation and classification

**Edge-weight validation against CollecTRI.** We stratified edges by $\phi =$ score_ctx_with_backfill into four tiers (with the overall distribution of non-zero $\phi$ shown in **Figure 5d**) and evaluated overlap with the CollecTRI manually curated gold standard[13]—an orthogonal resource not used in scaffold construction. In both states, CollecTRI overlap increased monotonically with $\phi$: Q4 (top quartile) edges showed 2.0–2.2$\times$ higher CollecTRI representation than Q1 (bottom quartile), confirming that $\phi$ distinguishes biologically validated regulatory relationships from spurious edges. Cross-state co-expression analysis provided further validation: ESC-specific edges ($\phi_{\text{ESC}} > 0.01$, $\phi_{\text{NPC}} \leq 0.01$) showed strong TF–target co-expression in ESC (mean Spearman $\rho > 0$) but negligible correlation in NPC, while NPC-specific edges displayed the opposite pattern. This confirms that the scaffold captures state-specific regulatory relationships rather than constitutive TF–target associations.

**Two-state edge classification.** Classifying all 303,769 pairs at $\tau = 0.01$ revealed NPC-specific edges (99,557; 43.6

#### Supplementary Note 7: Downstream cross-task transfer boundaries (sci-Plex and GDSC2)

**Cross-task boundary characterization.** To evaluate the boundaries of the framework, we tested VitaGRN on downstream tasks involving chemical/pharmacological perturbations rather than genetic (transcription factor) perturbations: the sci-Plex chemical perturbation dataset and the GDSC2 drug response screens. On these datasets, VitaGRN achieves competitive performance comparable to standard expression-based baselines, though it does not establish a new state-of-the-art predictive advantage (**Extended Data Fig. 7**). This comparable performance is consistent with the nature of small-molecule drug actions, which typically trigger complex, multi-target signaling cascades. While genetic perturbations directly disrupt specific transcription factors, chemical interventions involve broader pathway rewiring that partially diverges from the direct physical regulatory paths modeled by the sequence and structure engines. These analyses demonstrate that VitaGRN remains robust and competitive on chemical datasets, while highlighting that its unique biophysical advantages are most pronounced in genetic perturbation scenarios.

#### Supplementary Note 8: Detailed molecular mechanism and literature support for non-canonical translation factor EIF2B2

EIF2B2 encodes the beta subunit of eukaryotic translation initiation factor 2B (eIF2B), a decameric protein complex that serves as the guanine nucleotide exchange factor (GEF) for eIF2. Under physiological conditions, eIF2B catalyzes the exchange of GDP for GTP on eIF2, allowing the formation of the eIF2-GTP-Met-tRNA$_i$ ternary complex required for translation initiation. Knockdown or genetic mutation of EIF2B2 impairs this GEF activity, reducing the availability of the ternary complex and thereby attenuating global protein synthesis.

This reduction in functional eIF2B activity mimics the cellular integrated stress response (ISR) normally triggered by eIF2$\alpha$ phosphorylation. Paradoxically, while global translation is inhibited under these conditions, specific mRNAs containing inhibitory upstream open reading frames (uORFs) in their 5$'$ untranslated regions (UTRs) are translated more efficiently. The core ISR transcription factor ATF4 is the classic example of this translational control mechanism: under stress, delayed re-initiation allows ribosomes to bypass inhibitory uORFs and initiate translation at the downstream coding sequence of ATF4[15,16].

Once translated, ATF4 translocates to the nucleus and dimerizes with other bZIP transcription factors to activate genes containing amino acid response elements (AAREs) or unfolded protein response elements. Key target genes include DDIT3 (CHOP) and TRIB3, which mediate cell death and feedback pathways, and DDIT4 (REDD1), which acts as an inhibitor of mTORC1 signaling to conserve energy. Additionally, ATF4 directly activates the transcription of PHGDH, the rate-limiting enzyme in the phosphorylated pathway of serine biosynthesis, to sustain intracellular amino acid pools under stress[17,18]. The chaperone HSPA8 (HSC70) is also upregulated as part of the broader protein folding machinery activated during cellular stress. This indirect physical-to-transcriptional cascade (EIF2B2 $\rightarrow$ eIF2B complex GEF activity $\rightarrow$ global translation attenuation & selective ATF4 translation $\rightarrow$ transcriptional activation of ISR effector genes) is captured by VitaGRN through the PPI cascade, which maps physical interactions to trace non-canonical paths.

### Supplementary Tables

**Supplementary Table 1 | Primary K562 GRN benchmark results (Main candidate space $\mathcal{S}_{\text{main}}$ evaluated against reference standard $\mathcal{G}_{\text{ref}}$).** Methods are listed in descending macro-averaged AUPRC order; the proposed model is shown as **VitaGRN**.

| Method | AUPRC | Lift | AUROC | Recall | EP@50 |
|---|---|---|---|---|---|
| **VitaGRN** | 0.1342 | 32.32 | 0.6286 | 0.0748 | 0.1415 |
| scPRINT | 0.0937 | 22.57 | 0.5150 | 0.0026 | 0.0765 |
| PORTIA | 0.0926 | 22.31 | 0.5032 | 0.0263 | 0.0688 |
| scGNN | 0.0920 | 22.16 | 0.5081 | 0.0008 | 0.0044 |
| PSGRN | 0.0919 | 22.12 | 0.5099 | 0.0419 | 0.0840 |
| SpearmanCLR | 0.0913 | 21.98 | 0.5040 | 0.0029 | 0.0762 |
| GENIE3 | 0.0912 | 21.97 | 0.5100 | 0.0019 | 0.0044 |
| WGCNA-TOM | 0.0911 | 21.94 | 0.5092 | 0.0026 | 0.0653 |
| DAZZLE | 0.0910 | 21.91 | 0.5035 | 0.0006 | 0.0341 |
| PC | 0.0909 | 21.90 | 0.5023 | 0.0005 | 0.0055 |
| Geneformer | 0.0908 | 21.87 | 0.5015 | 0.0496 | 0.0764 |
| DeepRIG | 0.0907 | 21.83 | 0.5011 | 0.0002 | 0.0324 |
| PIDC | 0.0904 | 21.78 | 0.5057 | 0.0024 | 0.0691 |
| DeepSEM | 0.0904 | 21.77 | 0.5034 | 0.0010 | 0.0663 |
| CellOracle | 0.0904 | 21.77 | 0.5024 | 0.0004 | 0.0067 |
| Inferelator | 0.0903 | 21.74 | 0.5024 | 0.0003 | 0.0262 |
| SCODE | 0.0903 | 21.74 | 0.5015 | 0.0058 | 0.0422 |
| TENET | 0.0901 | 21.70 | 0.4995 | 0.0025 | 0.0524 |
| pySCENIC | 0.0898 | 21.63 | 0.5021 | 0.0004 | 0.0037 |
| GRNBoost2 | 0.0898 | 21.62 | 0.5006 | 0.0006 | 0.0032 |
| scGPT | 0.0896 | 21.56 | 0.4982 | 0.0312 | 0.0758 |

*Block scale:* 5,396 genes, 1,078 candidate TFs, 24,151 projected gold edges, and 50 evaluable gold TFs.

**Supplementary Table 2 | Perturbation response prediction benchmark across 5 methods and 6 metrics.** All values are averaged across the 218 held-out test transcription factors (217 for GEARS). VitaGRN achieves the best performance on all evaluated metrics.

| Method | PCC-$\delta$ | DEG20-PCC | DEG50-PCC | DirMatch | LFCSpear | RMSE-$\delta$ |
|---|---|---|---|---|---|---|
| **VitaGRN** | 0.495 | 0.681 | 0.686 | 0.635 | 0.370 | 0.076 |
| Matching Mean | 0.370 | 0.544 | 0.559 | 0.598 | 0.279 | 0.080 |
| GEARS | 0.339 | 0.553 | 0.561 | 0.583 | 0.220 | 0.086 |
| biolord | 0.298 | 0.462 | 0.475 | 0.566 | 0.180 | 0.081 |
| Latent Additive | 0.373 | 0.557 | 0.570 | 0.597 | 0.279 | 0.080 |

**Supplementary Table 3 | Complete extended biological case studies.** Top20 coverage: number of true top-20 DEGs covered by the VitaGRN local network. DirMatch@20: fraction of covered genes with correct predicted perturbation direction (denominator = coverage count, not 20).

| TF | Biological theme | PCC-$\delta$ | Top20 coverage | DirMatch@20 |
|---|---|---|---|---|
| EIF2B2 | eIF2 translation initiation / integrated stress response | 0.950 | 12/20 | 100% |
| DNTTIP2 | chromatin/RNA-associated nucleolar process | 0.783 | 11/20 | 100% |
| RPS6 | ribosomal protein / translation and growth signaling | 0.864 | 8/20 | 100% |
| DIMT1 | ribosome biogenesis / rRNA methylation | 0.820 | 13/20 | 100% |
| CDC6 | DNA replication licensing / cell cycle | 0.450 | 6/20 | 75% |
| DHX37 | RNA helicase / ribosome biogenesis | 0.760 | 14/20 | 100% |
| NUF2 | kinetochore / mitotic chromosome segregation | 0.832 | 12/20 | 95% |
| ETF1 | translation termination / protein synthesis | 0.808 | 13/20 | 100% |
| CHEK1 | DNA damage checkpoint / replication stress | 0.415 | 5/20 | 80% |
| CPSF3 | mRNA 3′ end processing / cleavage-polyadenylation | 0.460 | 4/20 | 95% |

**Supplementary Table 4 | Edge classification across ESC and NPC regulatory scaffolds.**

| Category | Edges | % of union |
|---|---:|---:|
| ESC-specific | 46,358 | 15.3% |
| NPC-specific | 99,557 | 32.8% |
| Shared-up (NPC) | 15,649 | 5.2% |
| Shared-down (ESC) | 30,439 | 10.0% |
| Shared-stable | 36,118 | 11.9% |
| Below threshold | 75,648 | 24.9% |

**Supplementary Table 5 | Top candidate TFs for experimental validation.** See Supplementary Methods for complete siRNA and qPCR protocols.

| **TF** | **Class** | **Direction** | **$\log_2$FC** | $\bar{\phi}_{\text{dom}}$ | **Disease relevance** |
|---|---|---|---|---|---|
| TP63 | Chromatin Rewirer | NPC | $-0.58$ | 0.788 | Neurodegeneration (Tau/TDP-43) |
| OVOL2 | Master Driver | ESC | $-9.59$ | 0.307 | Cancer EMT barrier |
| E2F7 | Chromatin Rewirer | NPC | $+0.74$ | 0.757 | ALS/FTD (FUS regulator) |
| IRF6 | Master Driver | ESC | $-10.37$ | 0.261 | Van der Woude syndrome |
| VDR | Master Driver | ESC | $-6.43$ | 0.245 | Multiple sclerosis (druggable) |
| ZNF410 | Chromatin Rewirer | NPC | $+1.12$ | 0.666 | Completely uncharacterized |

**Supplementary Table 6 | Hyperparameter search space and final configuration.** All selections based on validation set (175 TF) tuning.

| Parameter | Search range | Final | Selection criterion |
|---|---|---|---|
| SVD components $K$ | {10, 20, 30, 50, 100} | 20 | Validation PCC-$\delta$ saturates after $K = 20$; 61.7% variance retained |
| BayesianRidge max_iter | {100, 200, 300, 500} | 300 | Converges stably at 300; no gain beyond |
| Cosine neighbors $K_N$ | {3, 5, 10, 20} | 5 | Optimal validation DEG20-PCC at $K_N = 5$ |
| Cross-TF $\alpha_{\text{mix}}$ | {0.1, 0.3, 0.5, 0.7, 0.9} | 0.5 | Best from validation grid search |
| NN blend $\beta$ | {0.1, 0.2, 0.3, 0.5} | 0.3 | Weighted blend with Cross-TF output |
| ProTrek embedding dim | N/A | 1,024 | Fixed pre-trained model output |

**Supplementary Table 7 | Gold-GRN evidence strata, construction rules, and $\mathcal{S}_{\text{main}}$-projected scale.** All golds are intersected with the candidate space before scoring; $\mathcal{S}_{\text{main}}$ counts denote the positive edges and gold TFs actually used in scoring.

| Gold definition | Evidence source/rule | Role in this report | $\mathcal{S}_{\text{main}}$ gold edges | $\mathcal{S}_{\text{main}}$ gold TFs |
|---|---|---|---|---|
| $\mathcal{G}_{\text{ref}}$ | Integrated all-source physical-binding layer | Primary reference standard | 24,151 | 50 |
| CollecTRI | CollecTRI only (functional regulatory reference) | Single-source sensitivity | 233 | 30 |
| TRRUST | TRRUST only (literature regulatory reference) | Single-source sensitivity | 52 | 23 |
| BroadGRN | BroadGRN only (functional network reference) | Single-source sensitivity | 213 | 11 |
| ENCODE K562 | ENCODE K562 ChIP-seq (binding-only reference) | Binding-only sensitivity | 23,825 | 11 |
| Intersection $\geq 2$ | At least two source supports (strict intersection) | Small high-confidence audit | 37 | 13 |
| HiConf Expanded | Integrated functional and binding rules | Intersection robustness | 43 | 13 |
| Broad/ENCODE | Broad/ENCODE integration (expanded layer) | Extended audit | 24,151 | 50 |

**Supplementary Table 8 | K562 candidate-space definitions, scale, and intended use.** Candidate edges are directed TF–target combinations.

| Space | Role | Selection logic | Genes | TFs | Candidate edges | Limitation |
|---|---|---|---|---|---|---|
| Global ($\mathcal{S}_{\text{global}}$) | Coverage audit | All K562 genes and TFs | 5,968 | 5,966 | 35,605,088 | Very large TF background |
| Main ($\mathcal{S}_{\text{main}}$) | Primary space | Union of predicted edges in family | 5,396 | 1,078 | 5,816,888 | Restricted search domain |
| Shared ($\mathcal{S}_{\text{shared}}$) | Robustness space | Intersection of predicted edges in family | 3,293 | 978 | 3,220,554 | Narrowed denominator |

**Supplementary Table 9 | Complete data efficiency analysis (weighted cosine Cross-TF transfer).** Fixed 218 held-out TF test set; training TFs sub-sampled (3 seeds per fraction).

| Fraction | # TFs | Anchor PCC | +NN PCC | +GRN+NN PCC | GRN gain |
|---|---|---|---|---|---|
| 5% | 34 | 0.3880 | 0.3858 | 0.3951 | +0.0093 |
| 10% | 69 | 0.4026 | 0.3962 | 0.4178 | +0.0216 |
| 20% | 139 | 0.4127 | 0.4081 | 0.4325 | +0.0244 |
| 40% | 278 | 0.4360 | 0.4300 | 0.4604 | +0.0304 |
| 60% | 417 | 0.4458 | 0.4424 | 0.4752 | +0.0328 |
| 80% | 556 | 0.4502 | 0.4431 | 0.4799 | +0.0368 |
| 100% | 695 | 0.4501 | 0.4449 | 0.4801 | +0.0352 |

**Supplementary Table 10 | Complete structure distance quartile analysis** (218 test TFs).

| Quartile | # TFs | Mean distance | PCC-$\delta$ | DEG20-PCC | DirAcc |
|---|---|---|---|---|---|
| Q1 (close) | 55 | 0.216 | 0.617 | 0.797 | 0.904 |
| Q2 | 54 | 0.320 | 0.561 | 0.692 | 0.865 |
| Q3 | 54 | 0.377 | 0.422 | 0.637 | 0.800 |
| Q4 (far) | 55 | 0.437 | 0.380 | 0.596 | 0.782 |

**Supplementary Table 11 | Prediction quality stratified by perturbation effect magnitude.**

| Effect quartile | n TFs | Mean PCC-$\delta$ | Mean DEG20-PCC | Mean $\Delta$PCC (GRN benefit) |
|---|---|---|---|---|
| Q1 (smallest $\|\Delta\|$) | 55 | 0.416 | 0.651 | +0.011 |
| Q2 | 54 | 0.405 | 0.639 | +0.005 |
| Q3 | 54 | 0.462 | 0.699 | +0.020 |
| Q4 (largest $\|\Delta\|$) | 55 | 0.695 | 0.762 | +0.027 |

**Supplementary Table 12 | RPE1 cross-cell-line GRN topology evaluation ($\mathcal{S}_{\mathbf{main}}$ space).** VitaGRN uses zero-shot K562 GRN transfer without RPE1-specific training.

| Method | Tier 1 AUPRC | Tier 2 AUPRC | Tier 3 AUPRC | AUROC | Recall |
|---|---|---|---|---|---|
| VitaGRN | 0.0739 | 0.0407 | 0.0557 | 0.711 | 0.507 |

**Supplementary Table 13 | Prediction quality ranking of 218 held-out TFs (top 10 rows).**

| TF | PCC-$\delta$ | DEG20-PCC | Direction Acc. | ProTrek Distance |
|---|---|---|---|---|
| RPS23 | 0.904 | 0.995 | 1.000 | 0.380 (Q3) |
| RPL27 | 0.848 | 0.993 | 1.000 | 0.302 (Q2) |
| RPL21 | 0.933 | 0.990 | 1.000 | 0.270 (Q1) |
| DIMT1 | 0.819 | 0.989 | 1.000 | 0.388 (Q3) |
| RPL35 | 0.830 | 0.988 | 1.000 | 0.329 (Q2) |
| POLE | 0.857 | 0.988 | 0.984 | 0.158 (Q1) |
| EIF2B2 | 0.950 | 0.988 | 1.000 | 0.183 (Q1) |
| RPL19 | 0.930 | 0.988 | 1.000 | 0.311 (Q2) |
| RPS19 | 0.947 | 0.987 | 1.000 | 0.247 (Q1) |
| PSMC6 | 0.849 | 0.987 | 0.996 | 0.236 (Q1) |

**Supplementary Table 14 | Projected coverage of the reference standard ($\mathcal{G}_{\mathrm{ref}}$) and other gold layers across candidate spaces.** The table reports the positive scale actually scored after intersecting each gold with each space.

| Space | Gold layer | Genes | TFs | Projected gold edges | Evaluable gold TFs |
|---|---|---|---|---|---|
| $\mathcal{S}_{\mathrm{global}}$ | $\mathcal{G}_{\mathrm{ref}}$ | 5,968 | 5,966 | 230,769 | 405 |
| $\mathcal{S}_{\mathrm{main}}$ | $\mathcal{G}_{\mathrm{ref}}$ | 5,396 | 1,078 | 24,151 | 50 |
| $\mathcal{S}_{\mathrm{shared}}$ | $\mathcal{G}_{\mathrm{ref}}$ | 3,293 | 978 | 15,246 | 47 |
| $\mathcal{S}_{\mathrm{main}}$ | CollecTRI | 5,396 | 1,078 | 233 | 30 |
| $\mathcal{S}_{\mathrm{main}}$ | BroadGRN | 5,396 | 1,078 | 213 | 11 |
| $\mathcal{S}_{\mathrm{main}}$ | Intersect-ge2 | 5,396 | 1,078 | 37 | 13 |
| $\mathcal{S}_{\mathrm{main}}$ | HiConf-expanded | 5,396 | 1,078 | 43 | 13 |

**Supplementary Table 15 | Robustness summary for auxiliary gold definitions within the main candidate space $\mathcal{S}_{\text{main}}$.** All rows use the main candidate space $\mathcal{S}_{\text{main}}$ and are reported as evidence-sensitivity analysis.

| Gold definition | VitaGRN AUPRC | Best non-VitaGRN | Comparator AUPRC | Margin | Gold edges |
|---|---|---|---|---|---|
| $\mathcal{G}_{\text{ref}}$ | 0.134205 | scPRINT | 0.093729 | +0.040475 | 24,151 |
| CollecTRI | 0.073884 | scPRINT | 0.008234 | +0.065650 | 233 |
| BroadGRN | 0.040705 | scPRINT | 0.021813 | +0.018892 | 213 |
| TRRUST | 0.020515 | GENIE3 | 0.010383 | +0.010132 | 52 |
| ENCODE K562 | 0.403757 | PSGRN | 0.408798 | -0.005041 | 23,825 |
| Intersect-ge2 | 0.035278 | PC | 0.017155 | +0.018124 | 37 |
| HiConf-expanded | 0.035401 | GENIE3 | 0.012026 | +0.023375 | 43 |

**Supplementary Table 16 | Locally verified literature audit for evaluated model families.** Citation keys map directly to the expanded bibliography references in this study.

| Model or family | Role in the benchmark | Citation key |
|---|---|---|
| BEELINE | GRN benchmark design and metric context | [1] |
| GENIE3 / GRNBoost2 | Tree-based feature-importance GRN inference | [2] |
| CLR / SpearmanCLR | Mutual-information or correlation baselines | [3] |
| pySCENIC | Regulon inference and regulatory states | [5] |
| WGCNA-TOM | Co-expression and topological-overlap networks | [4] |
| CellOracle | Network inference and in silico TF perturbation | [6] |
| Inferelator | Sparse-regression regulatory-network learning | [19] |
| Inferelator 3.0 | Scalable single-cell GRN inference | [19] |
| PORTIA | Precision-matrix-based GRN inference | [7] |
| scGNN | Graph-neural-network framework for single-cell analysis | [20] |
| scPRINT | Pretrained cell model for gene-network prediction | [8] |
| PSGRN | GRN inference from perturbational single-cell data | [9] |
| DAZZLE | Dropout-augmented single-cell GRN inference | [10] |

**Supplementary Table 17 | Overview of evaluation datasets and tasks.** K562 and RPE1 form the main evaluation axis (GRN topology + response prediction); GDSC and sci-Plex are used for cross-task boundary characterization.

| Dataset | Scale | GRN topology evaluation | Downstream task evaluation |
|---|---|---|---|
| K562 | 152K cells, 1,092 perturbations | Main evaluation (22-method comparison) | Response prediction |
| RPE1 | 248K cells, 2,393 perturbations | Cross-cell-line GRN generalization | Cross-cell-line response transfer |
| GDSC | 947 cell lines, 286 drugs | — | Drug response prediction |
| Srivatsan 2020 | 799K cells, 188 drugs, 3 cell lines | — | Chemical perturbation transcriptome transfer |

**Supplementary Table 18 | K562 benchmark denominator control (Global candidate space $\mathcal{S}_{\mathrm{global}}$ evaluated against reference standard $\mathcal{G}_{\mathrm{ref}}$).** This table uses the same $\mathcal{G}_{\mathrm{ref}}$ reference but expands the background to the unconstrained global candidate space $\mathcal{S}_{\mathrm{global}}$. Methods are listed in descending macro-averaged AUPRC order; the proposed model is shown as **VitaGRN**.

| Method | AUPRC | Lift | AUROC | Recall | EP@50 |
|---|---:|---:|---:|---:|---:|
| scPRINT | 0.1019 | 15.72 | 0.5155 | 0.0036 | 0.1007 |
| **VitaGRN** | 0.1010 | 15.59 | 0.5161 | 0.0078 | 0.0175 |
| Geneformer | 0.1006 | 15.52 | 0.5124 | 0.0503 | 0.1074 |
| scGNN | 0.0993 | 15.31 | 0.5094 | 0.0392 | 0.0775 |
| pySCENIC | 0.0991 | 15.29 | 0.5036 | 0.0024 | 0.0521 |
| SpearmanCLR | 0.0981 | 15.13 | 0.5039 | 0.0057 | 0.1100 |
| WGCNA-TOM | 0.0980 | 15.11 | 0.5063 | 0.0089 | 0.0762 |
| CellOracle | 0.0980 | 15.11 | 0.5043 | 0.0050 | 0.0894 |
| GRNBoost2 | 0.0979 | 15.10 | 0.5059 | 0.0088 | 0.0692 |
| GENIE3 | 0.0977 | 15.07 | 0.5061 | 0.0062 | 0.0489 |
| DeepSEM | 0.0975 | 15.04 | 0.5031 | 0.0037 | 0.1019 |
| PC | 0.0973 | 15.00 | 0.5017 | 0.0017 | 0.0523 |
| PIDC | 0.0968 | 14.94 | 0.5024 | 0.0057 | 0.0795 |
| DeepRIG | 0.0964 | 14.88 | 0.5006 | 0.0005 | 0.0294 |
| DAZZLE | 0.0962 | 14.84 | 0.5018 | 0.0011 | 0.0436 |
| SCODE | 0.0961 | 14.82 | 0.5012 | 0.0037 | 0.0560 |
| PORTIA | 0.0959 | 14.80 | 0.5005 | 0.0041 | 0.0118 |
| PSGRN | 0.0959 | 14.79 | 0.5016 | 0.0077 | 0.0138 |
| Inferelator | 0.0959 | 14.79 | 0.5013 | 0.0004 | 0.0210 |
| TENET | 0.0957 | 14.77 | 0.5006 | 0.0031 | 0.0260 |
| scGPT | 0.0955 | 14.73 | 0.4999 | 0.0043 | 0.0130 |

*Block scale:* 5,968 genes, 5,966 candidate TFs, 230,769 projected gold edges, and 405 evaluable gold TFs.

**Supplementary Table 19 | K562 robustness benchmark block (Shared candidate space $\mathcal{S}_{\text{shared}}$ evaluated against the augmented broad+binding gold standard).** This table evaluates the shared candidate space $\mathcal{S}_{\text{shared}}$ representing the intersection of predicted genes and TFs. Methods are listed in descending macro-averaged AUPRC order; the proposed model is shown as **VitaGRN**.

| Method | AUPRC | Lift | AUROC | Recall | EP@50 |
|---|---|---|---|---|---|
| **VitaGRN** | 0.1459 | 30.80 | 0.6303 | 0.1183 | 0.1505 |
| scPRINT | 0.1027 | 21.68 | 0.5107 | 0.0035 | 0.0868 |
| PORTIA | 0.1021 | 21.56 | 0.5038 | 0.0235 | 0.0728 |
| scGNN | 0.1016 | 21.46 | 0.5100 | 0.0013 | 0.0060 |
| SpearmanCLR | 0.1011 | 21.36 | 0.5042 | 0.0037 | 0.0935 |
| GENIE3 | 0.1009 | 21.30 | 0.5079 | 0.0028 | 0.0055 |
| PSGRN | 0.1007 | 21.28 | 0.5057 | 0.0443 | 0.0906 |
| WGCNA-TOM | 0.1007 | 21.28 | 0.5099 | 0.0037 | 0.0735 |
| PC | 0.1003 | 21.19 | 0.5026 | 0.0008 | 0.0073 |
| DAZZLE | 0.1003 | 21.18 | 0.5038 | 0.0009 | 0.0452 |
| Geneformer | 0.1000 | 21.11 | 0.5029 | 0.0473 | 0.0834 |
| DeepRIG | 0.0999 | 21.10 | 0.5012 | 0.0002 | 0.0142 |
| CellOracle | 0.0999 | 21.09 | 0.5027 | 0.0007 | 0.0097 |
| DeepSEM | 0.0998 | 21.08 | 0.5037 | 0.0013 | 0.0752 |
| PIDC | 0.0998 | 21.07 | 0.5063 | 0.0031 | 0.0814 |
| Inferelator | 0.0997 | 21.06 | 0.5026 | 0.0005 | 0.0309 |
| SCODE | 0.0996 | 21.03 | 0.5014 | 0.0062 | 0.0476 |
| TENET | 0.0993 | 20.97 | 0.5000 | 0.0032 | 0.0565 |
| GRNBoost2 | 0.0990 | 20.90 | 0.5013 | 0.0009 | 0.0043 |
| pySCENIC | 0.0990 | 20.90 | 0.5024 | 0.0006 | 0.0054 |
| scGPT | 0.0984 | 20.79 | 0.4982 | 0.0287 | 0.0779 |

*Block scale:* 3,293 genes, 978 candidate TFs, 15,246 projected gold edges, and 47 evaluable gold TFs.
